# High-resolution genomic ancestry reveals mobility in early medieval Europe

**DOI:** 10.1101/2024.03.15.585102

**Authors:** Leo Speidel, Marina Silva, Thomas Booth, Ben Raffield, Kyriaki Anastasiadou, Christopher Barrington, Anders Götherström, Peter Heather, Pontus Skoglund

**Author notes:** Correspondence (L.S.), (P.S.).

## Abstract

Ancient DNA has unlocked new genetic histories and shed light on archaeological and historical questions, but many known and unknown historical events have remained below detection thresholds because subtle ancestry changes are challenging to reconstruct. Methods based on sharing of haplotypes^1,2^ and rare variants^3,4^ can improve power, but are not explicitly temporal and have not been adopted in unbiased ancestry models. Here, we develop *Twigstats*, a new approach of time-stratified ancestry analysis that can improve statistical power by an order of magnitude by focusing on coalescences in recent times, while remaining unbiased by population-specific drift. We apply this framework to 1,151 available ancient genomes, focussing on northern and central Europe in the historical period, and show that it allows modelling of individual-level ancestry using preceding genomes and provides previously unavailable resolution to detect broader ancestry transformations. In the first half of the first millennium ∼1-500 CE (Common Era), we observe an expansion of Scandinavian-related ancestry across western, central, and southern Europe. However, in the second half of the millennium ∼500-1000 CE, ancestry patterns suggest the regional disappearance or substantial admixture of these ancestries in multiple regions. Within Scandinavia itself, we document a major ancestry influx by ∼800 CE, when a large proportion of Viking Age individuals carried ancestry from groups related to continental Europe. This primarily affected southern Scandinavia, and was differentially represented in the western and eastern directions of the wider Viking world. We infer detailed ancestry portraits integrated with historical, archaeological, and stable isotope evidence, documenting mobility at an individual level. Overall, our results are consistent with substantial mobility in Europe in the early historical period, and suggest that time-stratified ancestry analysis can provide a new lens for genetic history.

## Introduction

Ancient genome sequencing has revolutionised our ability to reconstruct expansions, migrations and admixture events in the ancient past and understand their impact on human genetic variation today. However, tracing history using ancestry has remained challenging, particularly in (proto-)historical periods where the richest comparative information from history and archaeology often exists. This is because ancestries in many geographic regions are often so similar as to be statistically indistinguishable. One example is northern and central Europe since the start of the Iron Age c.500 BCE, where many longstanding questions remain, such as the nature of large-scale patterns of human migration during the 4th-6th centuries CE, their impact on the Mediterranean world, and later patterns of human mobility during the Viking Age (c. 750-1050 CE).

Reconstructing genetic histories from ancient DNA (aDNA) data commonly utilises so-called *f*-statistics^5–9^. Their popularity is rooted in a number of favourable properties, such as allowing analyses of lower-quality aDNA data, relative robustness to ascertainment, and theoretical guarantees of unbiasedness including in the presence of population bottlenecks^7,10^. *F*-statistics-derived approaches, such as *qpAdm^8^*, are close to unique in enabling the unbiased fitting of admixture events involving multiple source groups, including identifying the number of such events and choosing the best fitting sources^8,10,11^. However, *f*-statistics have not always had sufficient power to reconstruct events involving closely related ancestries, despite increasing sample sizes^12,13^. While the pairwise genetic differentiation (*F*_ST_) between the commonly studied hunter-gatherer, early farmer, and steppe-pastoralist groups that shaped the ancestry of Stone- and Bronze Age Europe^8,11,14,15^ is between 5% and 9%^8,16^, the differentiation between Iron Age groups in central and northern Europe is an order of magnitude lower at *F*_ST_=0.1-0.7% (Supplementary Figure 1). Methods that identify haplotypes, or shared segments of DNA that are not broken down by recombination, have previously been shown to have more power than single-SNP markers, but this information has not been accessible in combination with the advantages of *f*-statistics^2,13,17,18^. Furthermore, the overwhelming majority of available ancient DNA is from capture of 1.2 million SNPs^19^ and no clear advantages have been demonstrated for analysis of the >50 million SNPs available with whole-genome shotgun data as of yet.

One class of methods that use haplotype information is full genealogical tree inference, which can now readily be applied to many thousands of modern and ancient whole genomes^20–25^. These have been successfully applied to boost the detection of positive selection^22,26–28^, population structure^21,23,25,29^, infer geographic locations of ancestors^24,30^, demography^21,22^, and mutation rate changes^21^, among others. Genealogical trees can be thought of as containing essentially full, time-resolved information about ancestry, including information typically captured by recent haplotype sharing or identity-by-descent. In contrast, rare variant ascertainment, haplotypes, or chromosome blocks can be thought of as subsets or summaries of the information available in genealogies.

Here, we propose a new approach which we refer to as ‘time-stratified ancestry analysis’ to boost the statistical power of *f*-statistics several-fold by utilising inferred genome-wide genealogies (Figure 1a). We apply our method to the genetic history of northern and central Europe from c.500 BCE-1000 CE, from the start of the Iron Age to the Viking Age, using 1151 previously published whole-genome shotgun sequences (Supplementary Figure 1). We construct a fine-scale map of relationships between individuals and groups to understand ancestry shifts and expansions, for example of Germanic- and Slavic-speakers, and mobility in the Viking world.

**Figure 1.**
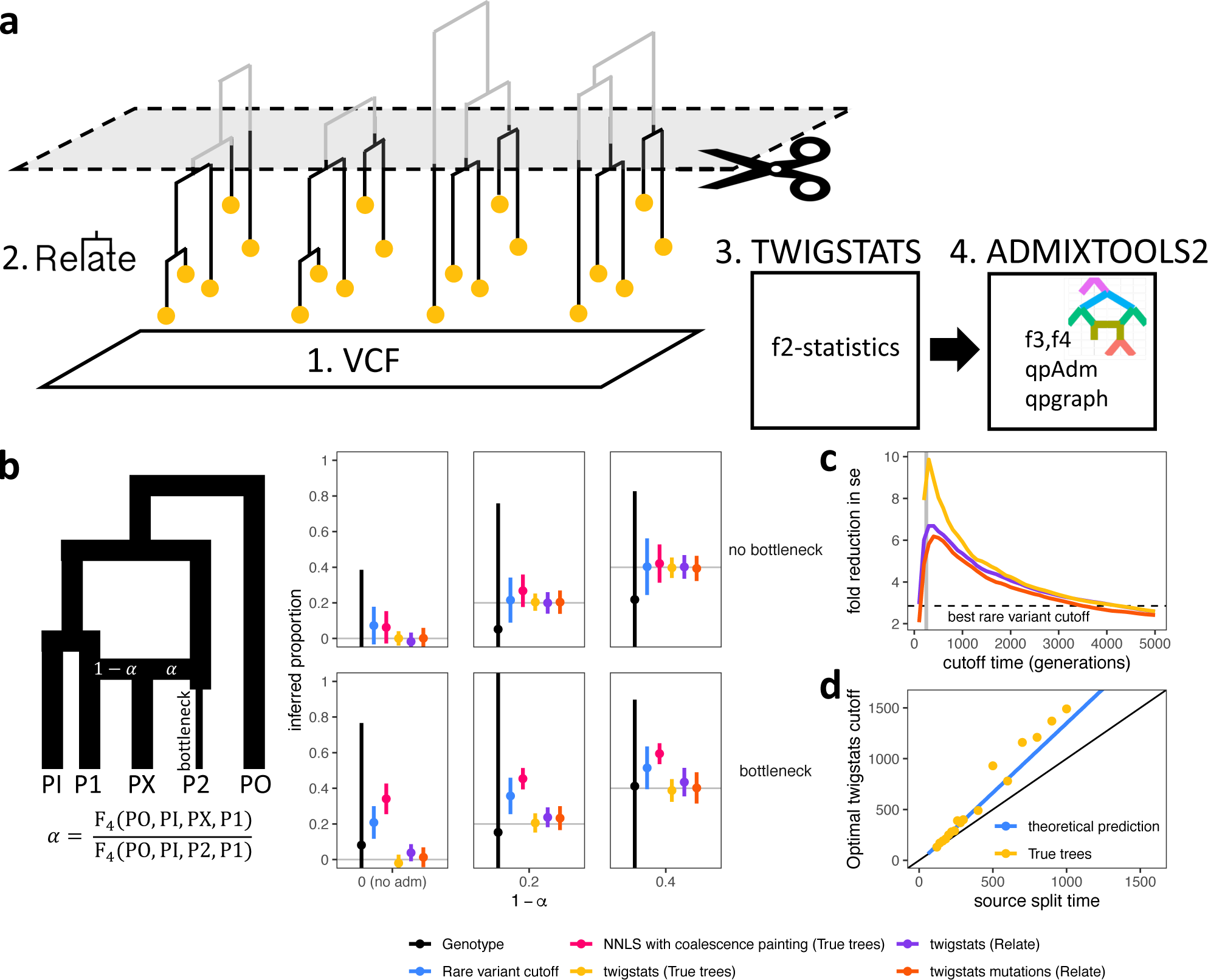
*Twigstats* performance on simulated data. **a,** Diagram of the *Twigstats* approach. We first construct genealogies from genetic variation data, and then use *Twigstats* to compute *f*_2_-statistics between pairs of groups to be used by ADMIXTOOLS2. **b,** Admixture proportions inferred from an *f*_4_-ratio statistic or non-negative least squares. Source groups P1 and P2 split 250 generations ago. Effective population sizes are equal and constant except for a recent bottleneck in P2 (see **Methods** for simulation details). The *Twigstats* cutoff is set to 500 generations, the rare variant cutoff is set to 5%, and we additionally infer admixture proportions by generating ‘first coalescence profiles’ for each population and modelling PX as a mixture of sources P1 and P2 using non-negative least squares (**Methods**). Two standard errors are shown. **c**, Fold improvement of standard errors relative to the genotype case as a function of the *Twigstats* cutoff time, for the same simulation as in b and averaged across different true admixture proportions. Dashed line shows the best standard error when ascertaining genotypes by frequency, when evaluated at different frequency cutoffs. **d,** Optimal *Twigstats* cutoff, defined as the largest reduction in standard errors relative to the genotype case, as a function of source split time in simulations using true trees. Blue line indicates our theoretical prediction (Supplementary Information).

## Results

### Genealogy-informed ancestry modelling improves power in simulations

*F*-statistics, by definition, count the occurrence of local genealogical relationships that are implied by how mutations are shared across individuals^31^. New genealogy inference methods trace common ancestors back in time beyond what is implied by individual mutations by utilising linkage information across genomic loci. By dating inferred coalescence events, genealogies add a time dimension onto genetic variation data; in particular, they enable, in principle, the best attainable age estimate of mutations from genetic variation^22,32^. This inherent relationship between *f*-statistics and local genealogies makes it straightforward to compute *f*-statistics directly on inferred genealogies^33^. Instead of computing *f*-statistics on observed mutations, they are now calculated on the inferred branches of genealogies, some of which may not be directly tagged by mutations but are inferred through resolving the local haplotype structure (**Methods**).

We develop mathematical theory and simulate a simple admixture model, in which the ancestry proportion is constrained in a single ratio of two *f*_4_-statistics^5^, to test this approach (Figure 1b, Supplementary Information). While unbiased, we find that using *f*-statistics computed on genealogies by itself does not yet yield a large improvement in statistical power for quantifying admixture events. However, we show, through theoretical prediction and simulation, that large improvements in power can be gained without bias by focusing on recent coalescences, which are most informative on recent admixture events (Supplementary Figure 3). Coalescences older than the time of divergence of the sources carry no information with respect to the admixture event and only add noise to the *f*-statistics. Excluding these therefore increases statistical power, while introducing no bias, in principle.

We implement this idea of studying the ‘twigs’ of gene trees in a new tool, *Twigstats* (Figure 1**a****, Methods**), which we demonstrate in simulations reduces standard errors by up to 10-fold and potentially more, depending on sample sizes and details of the genetic history model. The approach does not produce detectable bias in estimates of admixture proportions (Figure 1**b,c,d,** Supplementary Figure 3). Additionally, we demonstrate that computing *f*-statistics on mutation age ascertained genotypes produces a close to equal power gain to the use of full genealogies in many examples, while adding flexibility by allowing lower-quality genomes to be grafted onto a genealogy reconstructed with higher-quality genomes^21^.

Previous studies have suggested ascertaining rare mutations as a proxy for recent history^3,4^, but we show that this approach is prone to bias when effective population sizes vary, and that using full time-restricted genealogies is both unbiased and more powerful (Figure 1**b**, Supplementary Figure 3). We attribute this to the observation that mutation age is not always highly predictive of allele frequency (Supplementary Figure 2) and the genealogy-based approach gains power from the inclusion also of higher frequency young mutations that tag recent coalescences. We also demonstrate that a widely-used chromosome painting approach, where groups are modelled using a non-negative least squares of genome-wide painting profiles^2^, is also prone to bias, when source groups have undergone strong drift since the admixture event (Figure 1**b**, Supplementary Figure 3).

### Three empirical examples of applying *Twigstats* to study fine-scale ancestry

We next tested the *Twigstats* approach on a range of empirical examples. First, we used our time-restricted genealogy approach to boost pairwise outgroup *f*_3_-statistics^34^ to quantify fine-scale population structure, which we demonstrated using a previously proposed simulation^29^ (Supplementary Figure 4a). When applied to published genomes from Neolithic Europe, more than 4,500 years old, we were able to replicate previously suggested fine-scale structure between individuals buried in megalithic structures in Ireland compared to others^35^, which was apparent with time-restricted genealogies, but not with SNP data alone (Figure 2a).

**Figure 2.**
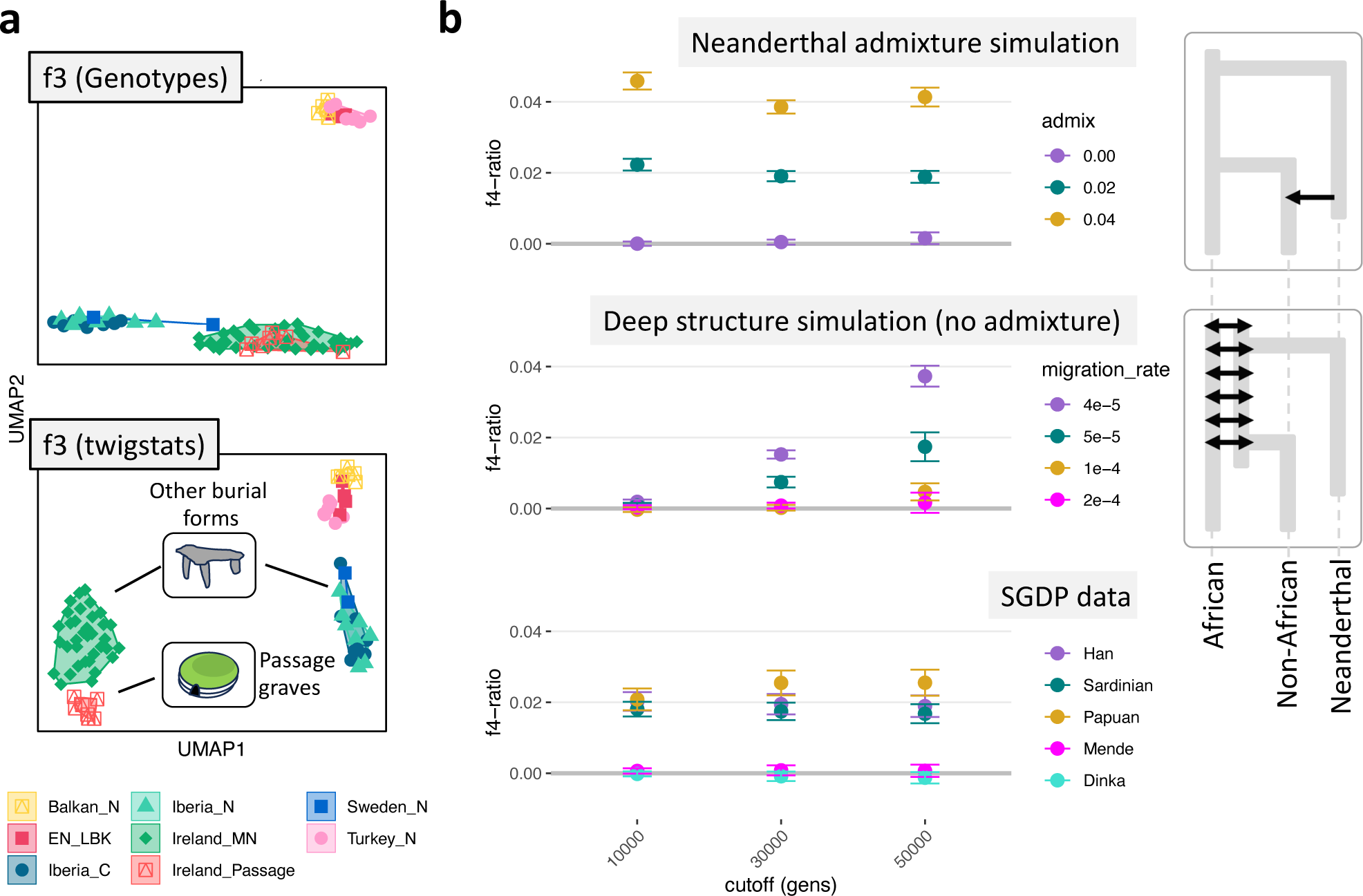
Two empirical applications where *Twigstats* time stratification reveals fine-scale structure and resolves timing of gene flow. **a,** Fine-scale genetic structure quantified using a *UMAP* calculated on a symmetric matrix containing pairwise outgroup *f*_3_ statistics (outgroup: YRI) between individuals calculated as conventional directly on genotype calls or calculated from *Relate* genealogies with a cutoff of 1000 generations. **b,** Neanderthal admixture proportion inferred using an *f*_4_-ratio of the form *f*_4_(outgroup, Altai, target, Mbuti)/*f*_4_(outgroup, Altai, Vindija, Mbuti). We compute equivalent *f*_4_-ratio statistics in a simulation emulating Neanderthal admixture 50,000 years ago and a second simulation involving no Neanderthal admixture but deep structure that leads to a similar inference unless deep coalescences are ignored by *Twigstats*.

Next, we applied the *Twigstats* approach to the well-studied example of three major ancestries contributing to prehistoric Europe: Mesolithic hunter-gatherers, early farmers, and steppe populations^8,11,14,15^. We obtain unbiased estimates and a ∼20% improvement in standard errors in an already well-powered *qpAdm* model for estimating these ancestries^36^ (Supplementary Figure 4c) and a remarkable power gain in distinguishing ancestries in Iron Age Northern Europe (Figure 3), mirroring our observation in simulations and theoretical prediction that the boost is most effective if differences between source groups are subtle (Supplementary Figure 3). For example, in models of admixture of British Iron Age and continental northern European ancestries in early medieval England^37^ (see below; Figure 3), *Twigstats* time-stratification halves standard errors from 8.0% with SNPs to 3.7% when using time-stratification (point estimates 65% and 73% Iron Age Britain-related ancestry, respectively)

**Figure 3.**
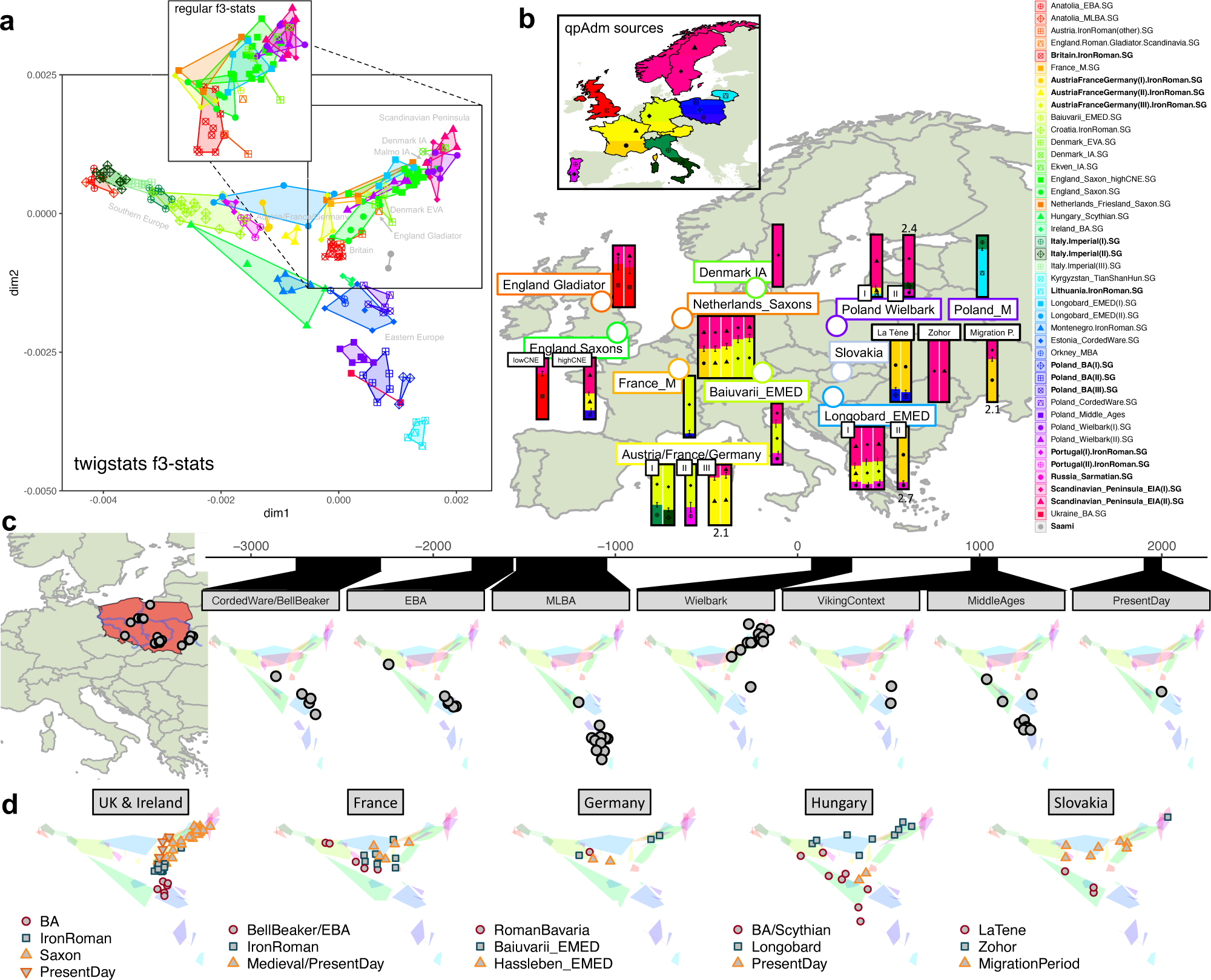
Ancestry in Iron Age and early medieval periods. **a**, Genetic structure of ancient groups from the Bronze Age to the medieval period quantified using a MDS of pairwise outgroup *f*_3_ statistics (outgroup: Han Chinese). These are calculated using *Twigstats* on Relate genealogies with a cutoff of 1000 generations. The magnified inset shows the MDS computed directly on genotypes. **b,** Ancestry models of Early Medieval groups across Europe computed using *qpAdm*. Sources are chosen from groups highlighted in the map and marked as bold in the legend, and were used in a rotational *qpAdm* scheme. For each target group, we remove models with infeasible admixture proportions and use a *Twigstats* cutoff of 1000 generations. All models satisfy p > 0.01, unless a -log10 p-value is shown next to the model. **c,** Ancestry change visualised over a time transect spanning 5000 years in present-day Poland (grey circles) plotted on the same MDS as in **a**. **d,** Ancestry change around the Iron Age and early medieval periods in the UK and Ireland, France, Germany, Hungary, and Slovakia plotted on the same MDS as in **a.**

Finally, we demonstrate that *Twigstats* can be used to resolve competing models of punctual admixture and long-standing gene flow, or constrain the time of admixture. For instance, it has previously been suggested that long-standing deep structure and gene flow between Neanderthals and early modern humans may produce patterns resembling a punctual admixture event some 60 thousand years ago^38–40^, and that some commonly used population genetic methods cannot distinguish between them, casting doubt on the model of Neanderthal admixture. However, while long-standing deep substructure in Africa does confound SNP-based *f*-statistics to produce patterns similar to Neanderthal admixture, we demonstrate, in simulations, that *Twigstats* can clearly distinguish this history from recent admixture (Figure 2**b**, Supplementary Figure 4b). Application of Twigstats on empirical whole genomes produces results inconsistent with deep substructure alone, but consistent with Neanderthal admixture into ancestors of Eurasians^40–42^.

### A framework for high resolution ancestry modelling in Metal Age and Medieval Northern Europe

Roman historians write extensively about the geographic distribution and extensive movements of groups and tribes immediately beyond the imperial frontier. The fall of the western Roman Empire has long been argued to have been both facilitated and in part caused by such population movements during the ‘Migration Period’. However, the exact nature and scale of these historically attested demographic phenomena—and their genetic impact—have been questioned due to uncertain historical sources and have been difficult to test with genetic approaches due to the close relations shared between many groups that were ostensibly involved. Less is understood at further distances from the Roman frontier due to lack of historical accounts.

Several recent studies have documented substantial mobility and genetic diversity in these time periods, suggesting stable population structure despite high mobility^43^, and documenting genetic variation in Viking Age Scandinavia^13,44,45^, early medieval England^3,37^, early medieval Hungary^46,47^, and Iron Age and medieval Poland^48^, but largely had to resort to using high-sample size modern cohorts to build models of ancestry change through time and space. Modern populations provide ample power to detect differences, but their genetic affinity to ancient individuals may be confounded by later gene flow, *i*.*e*. after the time of the ancient individual(s)^49^. The most principled approach is thus to build ancestry models where source and ‘outgroup/reference’ populations are older than, or at least contemporary with, the target genome or group which we are trying to model^49^. The improved statistical power of time-restricted ancestry in *Twigstats* thus offers an opportunity to develop principled ancestry models based solely on temporally appropriate source proxy groups.

To develop an ancestry model with sources from the Early Iron Age, we used hierarchical UPGMA clustering based on pairwise clade testing between all individuals, and formally tested the cladality of proposed ancestry groups with *qpWave* tests^43^ (**Methods**). This resulted in a set of model ancestry sources that included Iron Age and Roman Britain, the Iron Age of central European regions of Germany, Austria, and France, Roman Portugal, Roman Italy, Iron Age Lithuania, the Early Iron Age Scandinavian Peninsula (Sweden and Norway), as well as Bronze- and Iron Age Eastern and Southern Europe (Figure 3a,b, Supplementary Figure 1). Our source groups capture regional fine-scale genetic structure, such as separation of predominantly Norwegian and northern Swedish Early Iron Age individuals from southern Peninsular Scandinavia (Figure 3a), that is not detected without *Twigstats*. We employ a rotational *qpAdm* approach to narrow down the set of contributing sources from a larger pool of putative sources^50^. In addition, we made qualitative observations on results from non-parametric multi-dimensional scaling (MDS) on ‘outgroup-*f*_3_’ statistics^34^ (Supplementary Table 2).

### Expansion of ancestry related to Early Iron Age Scandinavia before and during the Migration Period

We assembled time transects using available ancient DNA data across several geographic regions in Europe, and inferred their ancestry using a model with Early Iron or Roman Age sources. Our modelling provides direct evidence of individuals with ancestry originating in northern Germany or Scandinavia appearing in these regions as early as the first century CE (Figure 3b, Supplementary Table 3).

In the region of present-day Poland, our analysis suggests several clear shifts in ancestry. First, in the Middle to Late Bronze Age (1500 BC - 1000 BC), we observe a clear shift away from preceding ancestry related to Corded Ware cultures^51^ ∼500CE. Second, in the 1st to 5th century CE, individuals associated with Wielbark culture^43,48^ represent an additional drastic qualitative shift away from the preceding Bronze Age groups, and can only be modelled with a >75% component attributed to the Early Iron Age (EIA) Scandinavian Peninsula, with multiple individuals at ∼100% and a stronger affinity in earlier Wielbark individuals. The Wielbark archaeological complex has been associated with the Goths, a Germanic-speaking group, but this attribution has remained unclear. Our modelling supports the idea that some early Germanic-speaking groups expanded into the area between the Oder and Vistula rivers, but since a considerable proportion of burials during this period were cremations, the possible presence of individuals with other ancestries can not be strictly rejected if they were exclusively cremated (and therefore invisible in the aDNA record).

A previous study could not reject continuity in ancestry from the Wielbark-associated individuals to later medieval individuals^48^. With *Twigstats’* improved power, models of continuity are strongly rejected, with no 1-source model of any preceding Iron Age or Bronze Age group providing a reasonable fit (p << 1e-33). Instead, individuals from medieval Poland can only be modelled as a mixture of majority Lithuanian Roman Iron Age ancestry, which is similar to ancestries of individuals from middle to late Bronze Age Poland, and a minority (95% CI: 18.9%-26.0%) ancestry component related to individuals from Roman Italy (p = 0.015) (Figure 3b). These medieval individuals from Poland thus carry no detectable Scandinavian-related ancestry, unlike the earlier Wielbark-associated individuals. Instead, present-day people from Poland are similar in ancestry to these medieval individuals (Figure 3c). The presence of a southern European-like ancestry component (represented by Roman Italy), which is not present in the Lithuanian Roman Iron Age, could be consistent with models of admixture taken place further south and arriving in Poland through north-westerly Slavic expansions.

In present-day Slovakia, individuals associated with the Iron Age La Tène period appear close to Hungarian Scythians in the two dimensions of our MDS analysis and are modelled as a mixture of central and eastern European ancestry. However, a 1^st^ century burial of a female aged 50-60 years from Zohor is modelled only with Scandinavian-related ancestry, corroborating the idea that this individual was associated with a nearby settlement that has been attributed culturally with Germanic-speakers^43,52^. Later Migration period individuals only have partial Scandinavian-related ancestry, suggesting later broader ancestry shifts in the region.

Nearby, in present-day Hungary, we observe Scandinavian-related ancestry components in several Longobard burials (Longobard_EMED(I)) dating to the middle of the 5th century CE^46^ (Figure 3b). This is consistent with the original study, which reported affinity to present-day groups from northwestern Europe (GBR, CEU, and FIN in the 1000 Genomes Project)^46^, but which we can extend with higher-resolution using earlier genomes. The Longobards were dominated by Germanic-speakers, and our result is consistent with attestations that they originated in the area of present-day Northern Germany or Denmark. In contrast, several other Longobard-associated individuals from the region (Longobard_EMED(II)) show ancestry more closely related to continental Europe, putatively representing local ancestry prior to Langobardic influence. This is consistent with the notion that the Longobards of the fifth and sixth centuries headed a confederation of several different groups. Present-day populations of Hungary do not appear to derive their ancestry from early medieval Longobards, and are instead more similar to Scythian ancestry sources (Supplementary Figure 5), consistent with the impact of later arrivals of Avars, Magyars and other eastern groups^53^.

In southern Germany, the genetic ancestry of individuals from early medieval Bavaria likely associated with the Germanic-speaking *Baiuvarii^54^*cannot be modelled as deriving ancestry solely from earlier groups in Iron Age Germany. Our current best model indicates a mixture with ancestry derived from EIA Peninsular Scandinavia, suggesting a regional ancestry shift and expansion of Scandinavian-related ancestry (Figure 3b). A subset of individuals from Iron Age and Roman Austria, France, and Germany are also best modelled with a minority Scandinavian-derived ancestry component. This may reflect expansions southwards—towards the Roman frontier—of other Germanic-speaking populations originating in Northern Europe.

In southern Britain, the ancestries of Iron Age and Roman individuals form a tight cluster in our MDS analysis (Figure 3a), shifted relative to available preceding Bronze Age individuals from Ireland and Orkney, and adjacent to, but distinct from, available individuals in Iron Age and Roman central Europe (Austria, France and Germany). However, two burials from a Roman military fortress site in Austria (Klosterneuburg)^43^ carry ancestry which is currently indistinguishable from Iron Age/Roman populations of Britain, to the exclusion of other groups (qpWave cladality p-value = 0.6). At least one of these skeletons (R10657 - Verf. Fn. 806/10), a 25-35-year old male buried beneath a horse^13^, shows osteological traits possibly linked to intense equestrianism, hinting that these may be non-local Roman soldiers. While one option is that they had ancestry from Britain, an alternative is that currently unsampled populations from large areas of western continental Europe carried ancestries similar to what we see in Iron Age southern Britain.

We further identify that an earlier Roman individual (6DT3) dating to ∼2nd century CE from the purported gladiator or military cemetery at Driffield Terrace in York (Roman *Eboracum*), England^55^, who was previously identified broadly as an ancestry outlier^56^, specifically carried EIA Scandinavian Peninsula-related ancestry. This documents an earlier presence of Scandinavian-related ancestry in Britain than in previous studies that suggested substantial influx of individuals carrying ‘continental northern European’ (CNE) ancestry associated with ‘Anglo-Saxon migrations’ from the 5th century onwards^37^. Roman-period individuals and groups from Northern Europe are indeed recorded in Roman sources both as soldiers and as slave gladiators condemned to die in Roman arenas^57,58^.

### Ancestry change in Late Iron Age southern Scandinavia

In Scandinavia, we found evidence for broad homogeneity in the Early Iron Age (<500CE). Specifically, individuals from Denmark 100CE - 300CE were indistinguishable from contemporary people in the Scandinavian Peninsula. However, we observe a clear shift in genetic ancestry already in the 8th century (Late Iron Age/early Viking Age) in Zealand island (present-day Denmark) for which a 100% Early Iron Age ancestry model is rejected (p = 1.2e-11 using *Twigstats*; p = 5e-3 without).

This shift in ancestry persists among later Viking Age groups in Denmark, where all groups are modelled with varying proportions of ancestry related to ‘continental’ Iron Age groups from present-day Austria, France, or Germany (Figure 4a,b). A non-parametric MDS of Viking Age individuals suggests that variation between individuals forms a cline spanning from the EIA Scandinavian Peninsula into continental Europe (Figure 4c). The observation of a shift in ancestry in Denmark could not be confounded by potentially unknown gene flow into the Iron Age groups in Austria, France, and Germany used in the model, but such gene flow could affect the exact ancestry proportions (Figure 3b).

**Figure 4.**
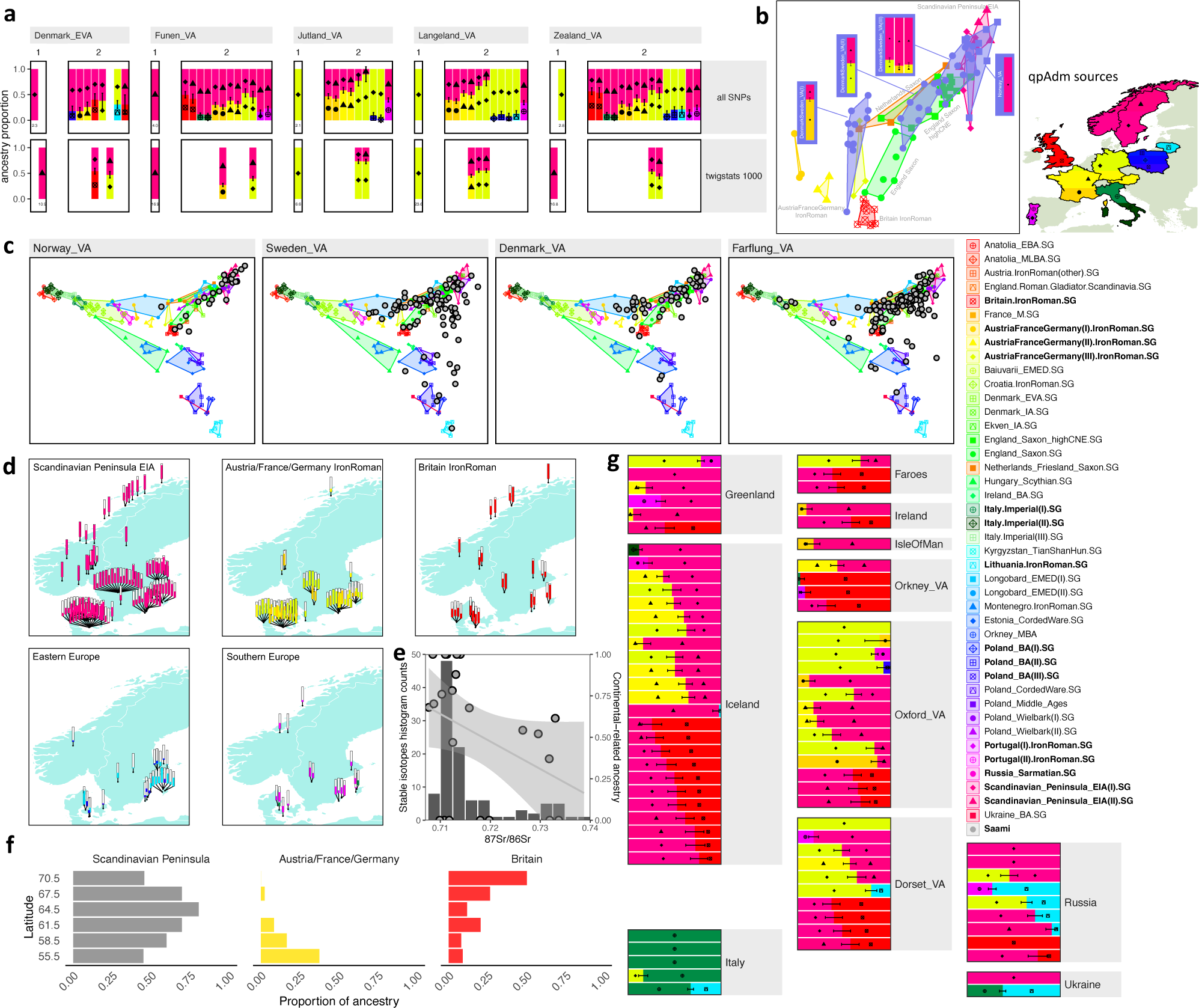
Ancestry in the Viking World. **a,** Ancestry shift observed in Viking Age Danish groups using *qpAdm* on all SNPs or *Twigstats*. We show the best 1 source and all accepted (p > 0.01) 2 source models. For rejected 1 source models, the -log10 p-value is shown under the plot. **b,** MDS and accepted *qpAdm* models (p > 0.01) for the main Danish, Swedish, and Norwegian Viking clusters identified with qpwave (Methods). **c,** Viking Age genetic variation (grey circles) visualised on the same MDS as in Figure 3a, but now including Viking age individuals. **d,** Map showing ancestry carried by Scandinavian Viking age individuals as inferred in their best-fitting *qpAdm* model. These are chosen by either choosing the 1-source model with largest p-value and p > 0.05 or the 2-source model with the largest p-value and p > 0.01. Supplementary Figure 6a shows the same map with all accepted models. **e,** Stable isotope data indicating the geology of childhood origin. Histogram shows the ratio of Strontium isotopes 87 to 86 measured in 109 individuals in Öland^68^. Grey circles indicate the correlation between continental ancestry and stable isotope values in a subset of these individuals (r = -0.39, p = 0.075). **f,** Ancestry proportion across Viking age individuals in panel **d** (Denmark, Sweden, and Norway) grouped by Latitude. **g,** Best fitting *qpAdm* ancestry model for farflung Viking individuals. Detailed models for all individuals are shown in Supplementary Figs 8 and 9.

These patterns are consistent with northwards gene flow into the Jutland peninsula and Zealand island in present-day Denmark, and expanding towards southern Sweden, potentially starting before the Viking Age. The timing is only constrained by the samples available, where the ancestry is not yet observed in the following latest individuals from south to north: individuals from Brøndsager and ⁄Kragehave Ødetofter in the Copenhagen area of Denmark dated to ∼100-300CE^13^, an individual from the southern tip of Sweden ∼500CE^15^, individuals from the Sandby Borg massacre site on Öland ∼500CE^44^, and a larger number of early Viking Age 8th century individuals from the Salme ship burial in present-day Estonia who likely originated in central Sweden^13^ (Supplementary Figure 8). Most likely the ancestry transformation thus occurred after those dates in each particular region, and mostly in the second half of the first millennium CE.

Interestingly, in southern Scandinavia, the archaeological record has yielded considerable evidence for periodic conflict and socio-political stress during the Iron Age, seen most explicitly in the many pre-Roman and Roman deposits of weapons and human remains in bogs^59^. While the context of these depositions remains debated and may have been variable, they could potentially relate to conflict associated with patterns of migration related to these ancestry shifts: a spread of other Germanic-speakers into the region from the south.

To assess the full extent of the impact of this ancestry influx into Scandinavia, we next turned to understanding the ancestry of individuals in Scandinavia during the Viking Age. Previous studies have suggested diversity of ancestries in Scandinavia in this period^13,44,60^, due to increased maritime mobility, but have not reported per-individual ancestry estimates based on preceding ancestry. We analysed each individual’s ancestry using a rotational *qpAdm* scheme (Figure 4d, Supplementary Table 4, Supplementary Figure 8), which showed increased power in distinguishing models when restricting to recent coalescences with *Twigstats* (>70% of accepted 1 source models in *Twigstats* were also accepted 1 source models using all SNPs, compared to <25% for the inverse). We were able to find a uniquely accepted model (p > 0.01) for 98 out of 165 individuals from Viking Age Scandinavia, compared to 70 using all SNPs, despite the fact that more *Twigstats* models had multiple ancestries (only 64 individuals with a 1 source model compared to 141 when using all SNPs) and were thus more complex.

We investigated regional differences in non-local ancestry across Scandinavia. In Denmark, 28 of 54 Viking Age individuals had detectable (z > 1) continental-related ancestry (Iron Age Germany, France or Austria) in their best accepted *qpAdm* models. In Sweden 28 of 74 individuals,, had detectable continental-related ancestry, concentrated almost entirely in southern regions (Figure 4d). By contrast, in Norway this ancestry was observed in only two individuals, indicating a wide-ranging impact of incoming ancestry in southern Scandinavia, and suggesting more continuity from the Early Iron Age in Norway and northern Sweden (Figure 4d). In more central parts of Scandinavia, the few individuals that show evidence of ‘continental’-related ancestry are observed in regions historically within the Danish sphere of influence and rule, such as the Oslo fjord. Relatively few such individuals are noted in eastern Central Sweden, which was a focus of regional power of the Svear (Figure 4d). The difference in distribution could suggest that the continental ancestry was more common in regions dominated by the historical Götar and those that lay on the borders of the Danish kingdom.

To test the extent to which the variation in ancestry was consistent with mobility during the lifetime of the individuals, or alternatively that of established groups, we focused on the island of Öland in southeast Sweden, where 23 individuals for which we could reconstruct ancestry portraits also have extensive associated strontium stable isotope data^61^. Strontium isotope data from dental enamel reflects the geology of the region where an individual grew to maturity, and there are considerable differences in expectations between Öland and many other regions in northern Europe. The full range of strontium isotope ratios in 109 individuals show two modes, a majority group with low ratios and a second minority group with high ratios falling outside the expected range of local fauna (Figure 4e).

Among 23 individuals with genomes in our data, all 5 individuals with clearly ∼100% continental ancestry (including one with ancestry related to Britain) are part of the majority strontium values, consistent with them having grown up locally. In contrast, the six most clearly non-local individuals based on the stable isotopes all have ∼50% or more EIA Scandinavian Peninsula-related ancestry, although five individuals with wholly EIA Scandinavian Peninsula-related ancestry also had local values. This suggests that the presence of continental ancestry was not a transient phenomenon, but an ancestry shift occurring at some point after the ∼500CE individuals from a massacre site at Sandby Borg ringfort on Öland, who all have strictly EIA Scandinavian-related ancestry.

### Origins of Viking Age mobility into Scandinavia

Previous studies had suggested a major influx of ancestry related to Britain into Viking Age Scandinavia^13,44^. While we detect this ancestry in some individuals (8 individuals in Norway, 12 in Denmark, 8 in Sweden), including some individuals whose ancestry appears to be entirely derived from Iron Age Britain, its overall impact appears reduced compared to previous reports. Our analysis indicates a proportionally larger impact of ancestry from Iron Age Britain in northern Norway, with southern Scandinavia predominantly influenced by continental ancestries (Figure 4f). While the sample sizes are limited, we observe nominal significance for an overrepresentation of females with British-related ancestry (Supplementary Figure 6c,d). We hypothesise that our estimates of ancestry from Britain are reduced relative to previous studies because the phenomenon of continental ancestry related to Iron Age France, Austria, and Germany may have been indistinguishable from ancestry from Britain. This could be due to a lack of statistical power to distinguish these closely related sources with standard methods, as well as through potential biases introduced by using modern surrogate populations that have since been influenced by later gene flow (such as gene flow into Britain). We illustrate this by replicating analyses of Refs ^13,44^ (Supplementary Figure 7).

Similarly, a previous study has suggested the presence of southern European ancestry at sites such as Kärda in southern Sweden^13^. In our models, two Kärda individuals are fit with continental ancestry, but none of the individuals have a substantial proportion of ancestry related to southern European sources (represented by Imperial Rome and Portugal in our analysis) (Supplementary Figure 7). Instead, southern European ancestry is only found in relatively small proportions in any individual in our models (Figure 4d).

Interestingly, we detect ancestry from Bronze and Iron Age sources from Eastern Europe (present-day Lithuania and Poland), concentrated in south-eastern parts of Sweden, particularly the island of Gotland (21 individuals, Figure 4d). This is consistent with previous genetic studies^13,44^. We find that this ancestry is enriched in males (Supplementary Figure 6d), suggesting male-biased mobility and/or burial. The closest match tends to be Roman Iron Age Lithuanian genomes associated with Balts, which would be consistent with mobility across the Baltic Sea, but we caution that the geographic representation of available genomes is still limited.

### Viking Age expansion from Scandinavia

Traditionally, historical perspectives on what is now often referred to as the ‘Viking diaspora’ placed an emphasis on the movements and settlements of population groups from various parts of Scandinavia^62^. Here, we use individual-level ancestry resolution to delve into the dynamics of Viking Age mobility, settlements, and interactions with local populations. When examining individuals in Viking-related contexts outside of Scandinavia, our explorative MDS analysis again indicates mixed ancestries related to the Scandinavian EIA, with regional differences that point to varied local admixture (Figure 4**c**, Supplementary Figure 9).

In Britain, most individuals recovered from the late Viking Age sites of Ridgway Hill, Dorset, and St. John’s College, Oxford^13^ show ancestries typical of those seen in Viking Age southern Scandinavia, suggesting that those executed could have been predominantly migrants or members of Viking raiding parties from Scandinavia (Figure 4g). In agreement with these genetic ancestry profiles, strontium and oxygen analyses of the individuals from Ridgway Hill and St. John’s College have suggested childhood origins outside of Britain, and potentially in peninsular Scandinavia, particularly the island of Öland^63,64^, or parts of northern continental Europe^65^. Notably, VK166 (SK 1891) from St. John’s College is a slight outlier in terms of both genetic ancestry, which is best explained by groups from Iron Age Austria, France, and Germany and Sarmatian-related populations from Russia, and the stable isotope analysis, where the oxygen results are suggest a more continental origin and the strontium suggests he spent his childhood on more radiogenic geology.

Further west, North Atlantic Viking Age individuals in the Faroe Islands, Iceland and Greenland show Scandinavian origins with several individuals showing the continental ancestry signal (Figure 4g). We observe that some individuals interred within pre-Christian burials in Viking Age Iceland share substantial ancestry with Iron Age Britain, which is consistent with previous studies^45^ and historical records. Previous hypotheses have included female-biased arrival of ancestry related to Britain and Ireland on Iceland^66^. We tested this but, contrary to this prior hypothesis, we found a marginal enrichment of ancestry related to Britain/Ireland/continental Europe in males (12 of 17 males and 2 of 7 females with at least one accepted model involving Iron/Roman Age Britain as source; Fisher’s exact test p = 0.085) (Supplementary Figure 6c). However, sampling of additional individuals that will improve distinction of Anglo-Saxon-related and Norse related-ancestries would be required to fully test this hypothesis.

In eastern Europe, we observe similar EIA Scandinavian ancestries in a Viking Age burial from Ukraine, and these ancestries are overrepresented in Viking Age burials from present-day Russia. At Staraya Ladoga in western Russia, we observe several individuals with EIA Scandinavian Peninsula-related ancestry and at least one individual dated to the 11th century with apparent ancestry related to Iron Age Britain. The relative absence of continental ancestry, which was largely restricted to southern Scandinavia during the Viking Age, is thus inconclusive but indicative that these individuals originated in the central/northern parts of Sweden or Norway.

## Conclusions

Our new approach, *Twigstats*, transfers the power advantage of haplotype-based approaches to a fully temporal framework, which is applicable to *f*-statistics and enables previously unavailable unbiased and time-stratified analyses of admixture. We demonstrated that *Twigstats* enables fine-scale quantitative modelling of ancestry proportions, revealing wide-ranging ancestry changes impacting Northern and Central Europe in the Iron, Roman, and Viking ages. We reveal evidence of expansion southwards and/or eastwards of likely Germanic speakers carrying Scandinavian-related ancestry in the first half of the first millennium CE. We note that ‘Scandinavian-related’ in this context relates to the ancient genomes available, and so it is entirely possible that these processes were driven e.g. from regions in northern-central Europe. This could be consistent with the attraction of the greater wealth, which tended to build up among Rome’s immediate neighbours and may have played a major role in vectors of migration internal to communities in Europe who lived beyond the Roman frontier^67^. Later, patterns of gene flow seems to have turned northwards, with the spread of continental-related ancestry in Scandinavia, which is later reflected in differential ancestries in the western and eastern directions of the Viking diaspora. Overall, our results suggest that whole-genome sequencing offers additional power over SNP capture through the use of time-restricted ancestry analysis, and opens up the possibility of the reconstruction of new high-resolution genetic histories around the world.

## Acknowledgements

L.S. was supported by a Sir Henry Wellcome Fellowship (220457/Z/20/Z). P.S. was supported by the European Molecular Biology Organisation, the Vallee Foundation, the European Research Council (grant no. 852558), the Wellcome Trust (217223/Z/19/Z), and Francis Crick Institute core funding (FC001595) from Cancer Research UK, the UK Medical Research Council, and the Wellcome Trust. B.R. was supported by the Swedish Research Council (grant no. 2021-03333).

## Code availability

*Twigstats* is freely available under an MIT licence through github https://github.com/leospeidel/twigstats and detailed documentation, as well as example data, is available under https://leospeidel.com/twigstats/.

## Data availability

All ancient DNA data used in this study was publically available and is listed in Supplementary Table 1.

## Author contributions

P.S. supervised the study. L.S. and P.S. developed the method, L.S, M.S., and P.S. curated the dataset, L.S. and P.S. analysed the data, all authors interpreted results and wrote the manuscript.

## Methods

### Twigstats

*Twigstats* takes the *Relate* output format as input and allows computation of *f*-statistics directly on genealogies, by using the inferred expected number of mutations on each branch as input, which is computed as the product of a prespecified average mutation rate per base per generation, the branch length, and the number of bases each tree persists^33^. Importantly, *Twigstats* computes *f*_2_-statistics ascertained by an upper date threshold, such that only branches younger than this threshold are used. If a branch crosses the threshold, we only use the proportion of the branch underneath the threshold. *Twigstats* additionally allows specifying a minimum derived allele frequency and lower date-threshold. *Twigstats* can also compute *f*_2_ statistics on age-ascertained mutations, which is particularly convenient for individuals not built into the genealogies.

The computed *f*_2_-statistics are fed into *admixtools2^69^*for computing derived statistics, which implements computation of genome-wide *f*_2_, *f*_3_, and *f*_4_ statistics, as well as *qpgraph* and *qpAdm* models. We implement the sample size correction as detailed in Ref ^7^. The *f*_2_ statistics are computed in blocks, typically of prespecified centi-Morgan size, or of prespecified physical distance. These blocks are used downstream within *admixtools2* to compute standard errors using a block-jacknife approach. By default, we compute *f*-statistics only on internal branches and exclude singleton tip branches to increase robustness against sample age.

The optimal *Twigstats* time cutoff is *a priori* unknown, however we develop theory that predicts the optimal choice in a simple 2-way admixture as a function of the admixture date, source split time, and admixture proportion (Supplementary Information). In this case, the optimal cutoff equals approximately 1.4 times the split time between admixing source groups, depending on exact parameters in the model (Figure 1**b,c**, Supplementary Figure 3).

### Non-negative least squares ancestry modelling

We implement an approach that uses genealogies to emulate the chromosome painting technique of identifying closest genetic relatives along the genome^1,2^ to fit admixture weights. When applied to true genealogies in simulations, this approach represents an idealised version of this idea.

Given known assignment of each sample to a population, we implement a function in the *Twigstats* R package that identifies, at each position in the genome, the population with which a sample coalescences first. Our implementation takes a list of reference populations as input, such that any coalescences not involving these reference populations are ignored when traversing back in time through genealogical trees. If the first coalescence involves multiple different reference populations, this coalescence event will be assigned to each population with weight proportional to the number of samples in each population involved in that event.

We then compute, for each target population and putative source populations, the proportion of the genome where the first coalescence involves each of the reference populations. Given *k* reference populations, we denote by *a_i_* the vector of length *k* storing these proportions for population *i*. We fit our target population as a mixture of putative source populations using a non-negative least squares approach that finds a solution to the following optimisation problem *min*_0≤Σβ_l_≤1_ || *a*_*t*_ − *Aβ* ||_2_, where *t* indexes the target population, *A* is a matrix storing *a_i_* for putative source populations as its column vectors and *β* are non-negative mixture weights.

### Admixture simulations

We use msprime^70^ to simulate genetic variation data to test our approach.

#### *F*_4_-ratio admixture simulation

Our simulation in Figure 1b and Supplementary Figure 3b,c simulates five populations named PI, PO, P1, P2, and PX, where PO splits from all other populations 10,000 generations ago, P1 and P2 represent two proxy source groups that split from each other at 250 generations or 500 generations ago, PI splits from P1 100 generations ago and PX emerges from a pulse admixture between P1 and P2 50 generations ago. All populations have a constant diploid population size of 5000 and a variable human-like recombination map, where our simulation only covers chromosome 1, and a human-like mutation rate of 1.25e-8 mutations per base per generation. We additionally have a modified simulation with a lower mutation rate of 4e-9 mutation per base per generation emulating a transversions-only data set and a simulation where P2 has a diploid population size of 1000 in the last 50 generations, emulating a recent bottleneck in this population. We sample 20 haploid sequences from all populations. The “large sample size” simulation samples 100 haploid sequences from all populations.

### *qpAdm* simulation

Our simulation in Supplementary Figure 3d uses the simulation model and script provided with ref. ^10^, where we changed this script to use the human hotspot recombination map. We simulated only chromosome 1. In the original simulation model, admixing sources split 1200 generations ago, with admixture occurring 40 generations ago. We additionally simulate a version where all population split times and admixture times are reduced by a factor of 5.

#### Neanderthal admixture and deep structure simulation

Our simulation in Figure 2b and Supplementary Figure 4c emulates Neanderthal admixture, where Neanderthals and ancestors of modern humans split 25,000 generations ago and admixture occurs 2,000 generations ago. The resulting admixed ‘non-African’-like population coexists with the non-admixed ‘African’-like population until the present-day. Additionally, two Neanderthal populations split from each other 7,000 generations ago, which can be interpreted as emulating the Altai and Vindija Neanderthal populations, with Vindija being closer to the admixing source.

Alternatively, we simulate a model with two subgroups emulating ancestral modern humans in Africa that have a non-zero symmetric migration rate, ranging from 4e-5 to 2e-4 per generation, up until 3,000 generations before present. One of these subgroups gives rise to a present-day ‘African’-like population, while the other gives rise to a present-day ‘non-African’-like population. We further sample two Neanderthal populations that split 7,000 generations ago and merge 25,000 generations ago with the same ancestral modern human subgroup that will eventually give rise to non-Africans.

We simulate whole genomes with human-like recombination rates and a mutation rate of 1.25e-8 mutations per base per generation. Diploid effective population sizes are set to 10,000 except on the Neanderthal lineage where it is set to 3,000. We sample 2 haploid sequences for each Neanderthal population and 20 haploid sequences for the target admixed population and African non-admixed population.

#### Fine-scale structure simulation

Our simulation in Supplementary Figure 4a emulates the emergence of fine-scale population structure and is adapted from ref. ^29^. In this simulation, populations split 100 generations ago into 25 sub-populations followed by a period in which individuals are allowed to migrate at a rate of 0.01 between adjacent populations in a 5x5 grid. The diploid effective population size is 500 in each of the 25 populations, and 10,000 in the ancestral population. We simulate 10 replicates of chromosome 10, with a human-like mutation rate of 1.25e-8 and hotspot recombination map. We sample 2 diploid individuals from each population. In addition, we sample 100 individuals from an ancestral population that splits from the 25 target populations 100 generations ago, prior to the emergence of structure in these 25 populations. *Relate* trees are inferred assuming true mutation rates, recombination rates, and average coalescence rates across all samples.

### Ancient sample selection

A full list of ancient genomes can be found in Supplementary Table 1. Published ancient shotgun genomes provided by Refs. ^44,45^ were only available aligned against GRCh38. These data were realigned to GRCh37d5 reference using *bwa aln* (version 0.7.17-r1188).

We select genomes with average autosomal coverage exceeding 0.5x with exception of VK518 of previously suggested to be of Saami ancestry who has coverage of 0.438 and was included to capture this ancestry in our panel. Genomes above a coverage cutoff of 0.5x have previously been shown to result in reliable imputation results^71^. We exclude samples with evidence of contamination. We remove any duplicate individuals, such as individuals that were resequenced, choosing the file with higher coverage. We then filter out any relatives annotated in the Allen Ancient DNA Resource v54.1^19^, retaining the individual with highest coverage in each family clade.

Our final dataset includes 1151 ancient genomes.

### Imputation of ancient genomes

We follow the recommended pipeline of *GLIMPSE^72^*, and first call genotype likelihoods for each genome at 1000 Genomes Project segregating sites using bcftools mpileup with filter -q 20, -Q 20, and -C 50. We subsequently impute each genome separately using GLIMPSE v1.1.1 using the 1000 Genomes Project Phase 3 reference panel^73^. These imputed genomes are merged into a single VCF for further downstream processing.

We filter any site where more than 2% of sites have an imputation posterior of less than 0.8 and retain all remaining sites so as to not have any missing genotypes at individual SNPs. We merge these ancient genomes with a subset of the 1000 Genomes Project data (Supplementary Table 1) to obtain a data set of a total of 1842 genomes.

#### *Relate*-inferred genealogies

We merge imputed ancient genomes with a subset of the 1000 Genomes Project dataset, including all European populations, CHB (Chinese Han from Beijing), and YRI (Yoruba). We create a second dataset where we merge imputed genomes with the Simon Genome Diversity Panel. We then infer genealogies for the joint data set of ancient and modern genomes using Relate v1.2.1. We restricted our analysis to transversions only and assume a mutation rate of 4e-9 mutations per base per generation and input sample dates as shown in Supplementary Table 1. We use coalescences rates pre-inferred for the 1000 Genomes project.

#### *QpAdm* modelling

Briefly, *qpAdm* models are a generalisation of *f*_4_-ratios, where 1-, 2-, and 3-source models can be tested as hypotheses, and admixture components and their standard errors obtained with a block jackknife^8^. A *qpAadm* model is fully specified by a set of putative source groups and additional ‘outgroups’ that are used to distinguish source ancestries. We employed a rotating approach where we iteratively selected a subset of source groups, and used all remaining putative sources as outgroups. This approach penalises models where true contributing sources are used as outgroups. With sufficient statistical power, *qpAdm* models will be statistically rejected if true contributing sources are used as outgroups. If statistical power is more limited, several models will fit the data, but the correct model is expected to be preferred over wrong models.

#### Clustering using *qpwave*

To identify homogeneous groups that we use as source groups in *qpAdm* modelling in Figures 3 and 4, we follow ref. ^43^ and run *qpwave* using *Twigstats* between pairs of ancient individuals. We use CHB and five European population from the 1000 Genomes Project as reference groups. This approach tests if two individuals form a clade with respect to reference groups. The reason why this is a principled approach despite the 1000 genomes groups postdating the ancient individuals is that if a group of ancient individuals are truly homogeneous, they will be so with respect also to postdating individuals.

We then define clusters by running UPGMA on -log10 p-values obtained from *qpwave* between all pairs of individuals and cut the resulting dendrogram at a height corresponding to a p-value of 0.01. We subsequently intersect cluster labels with pre-defined population labels.

To choose source groups for Figure 3, we run this algorithm on samples from Britain.IronRoman, AustriaFranceGermany.IronRoman, Scandinavian_Peninsula_EIA, Poland_BA, Lithuania.IronRoman, Portugal.IronRoman, and Italy.Imperial (Supplementary Table 1). We require a source group to have at least three individuals and therefore exclude clusters of size one or two.

This approach results in two clusters in the Scandinavian Peninsula, approximately separating northern from southern Scandinavia, three clusters in Poland that separate samples temporally between the early and later Bronze Age and two clusters for Iron and Roman Age Portugal. In present-day Austria, Germany, and France this approach identifies three clusters, with each cluster spanning multiple archaeological sites in different countries, indicating genetic diversity in this region in the first millenium CE. In Figure 4b, we run this algorithm on Viking Age samples from Norway, Sweden, and Denmark dating to <1000CE. To restrict to the main genetic clusters, we select all groups with at least five individuals. This results in three clusters of samples in Sweden and Denmark and a single cluster in Norway.

## Supplementary Figures

**Supplementary Figure 1.**
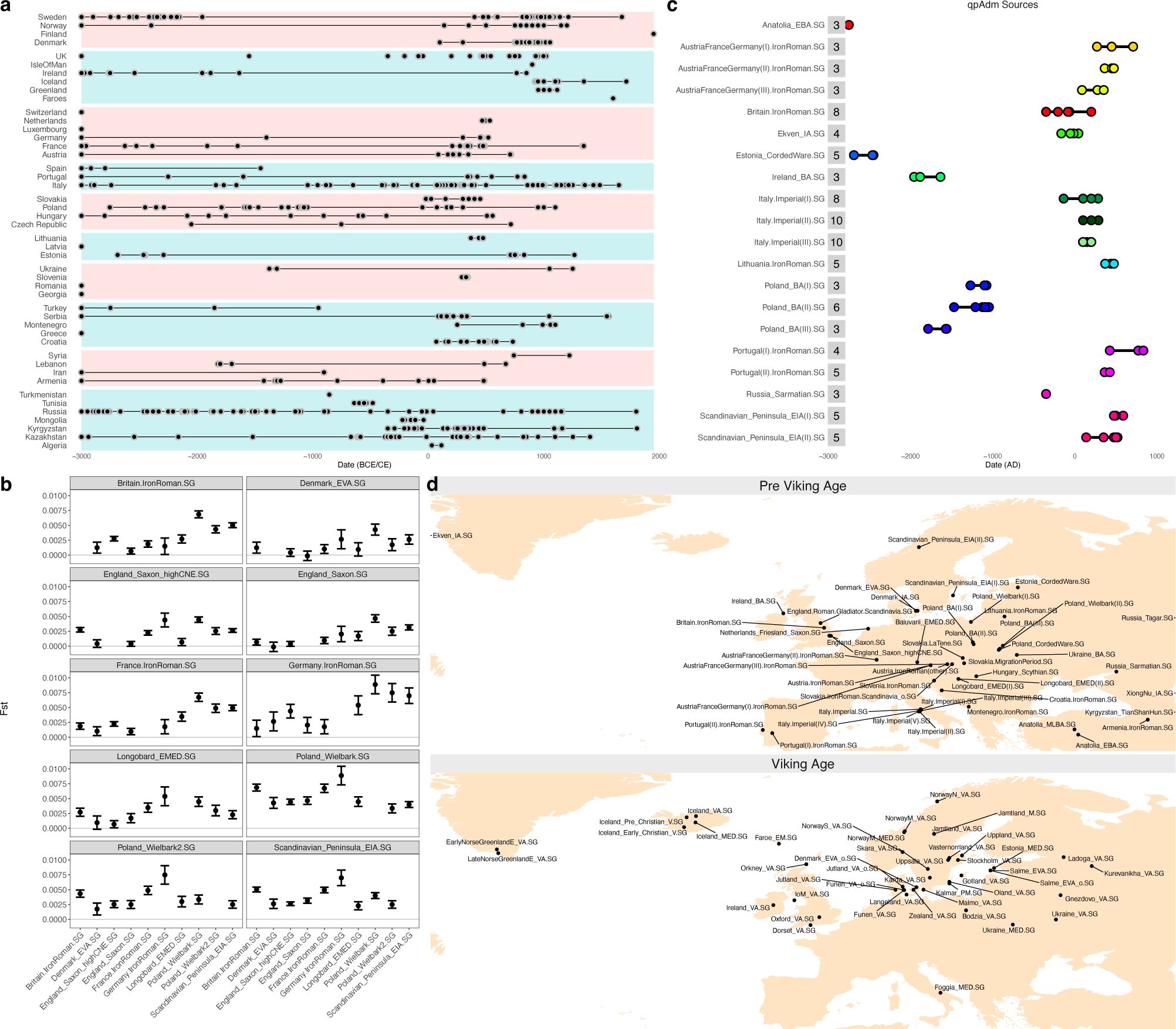
Collection of ancient genomes used in this study. **a,** Ancient DNA samples included in this study (Supplementary Table 1). Samples older than 3000 BCE are shown at 3000 BCE. **b,** Fst between Pre Viking Age groups computed using popstats^74^ using options --FST --informative. We show 1 standard error. **c,** Source groups used in *qpAdm* modelling of Metal Age and Medieval Northern Europe (Figures 3 and 4), showing sample sizes and sample ages. **d,** Map showing mean coordinates of sampled individuals in pre Viking Age and Viking Age groups.

**Supplementary Figure 2.**
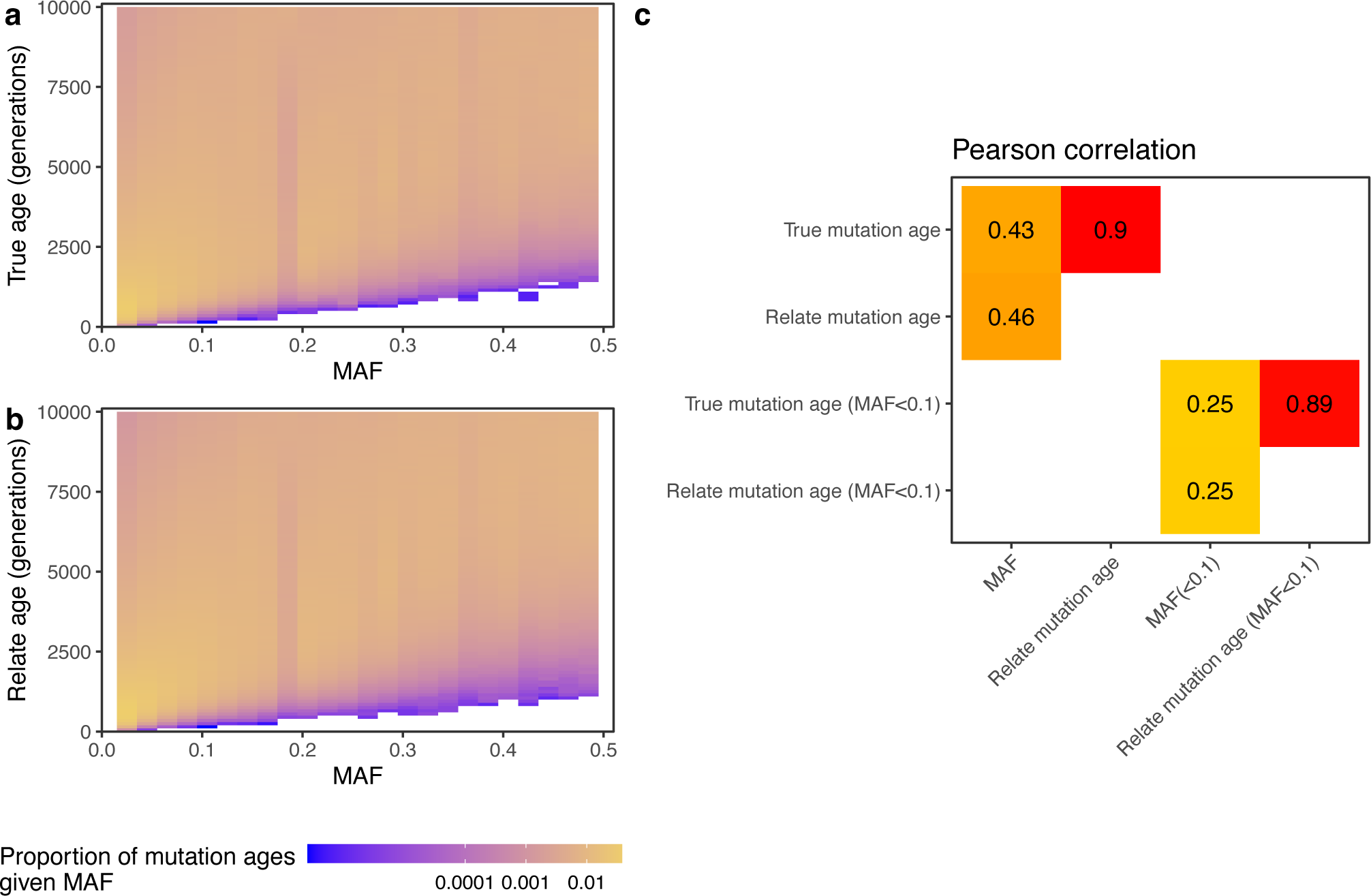
Relationship of mutation age and MAF. **a,** Heatmap showing the distribution of mutation ages for each minor allele frequency (MAF) bin. To account for the uncertainty in when the mutation arose on a branch, we sample a random date between the lower and upper ends of the branch onto which it maps. We use 20 replicates of the simulation of Fig 1b, where sources split 500 generations ago and the admixture proportion is 0%. **b,** Same as **a** but using mutation ages determined by *Relate*-inferred genealogies. We place mutations at the same relative height between the lower and upper ends of a branch as in the true trees to remove the uncertainty in when on the branch the mutation occurred, so that we would recover the true allele age from a correctly inferred genealogy. **c,** Pearson correlation between MAF, true mutation age, and Relate mutation ages, as well as the same comparisons when restricting to mutations of MAF less than 0.1.

**Supplementary Figure 3.**
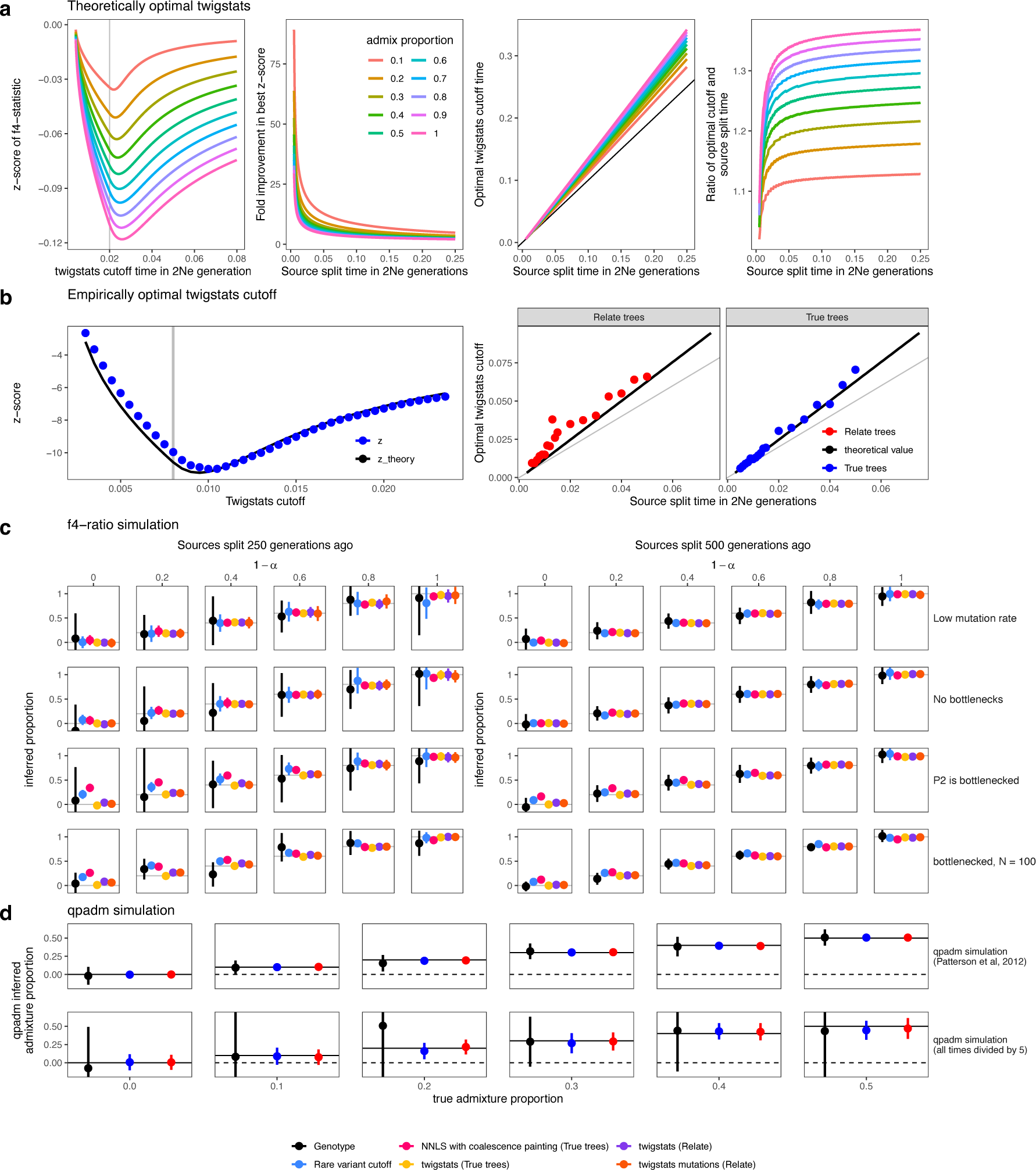
*Twigstats* optimal cutoff and simulation performance. **a,** Theoretically computed z-score of *f*_4_(PO,P1,PX,P2) at a single genomic locus (Supplementary Information), assuming PX is admixted between P1 and P2 at time 0.004 (in units of 2Ne generations), e.g. corresponding to 100 generations with 2Ne of 25,000. Sources split at time 0.02 in the first panel. We show how the optimal z-score and *Twigstats* cutoff behaves as a function of the source split time and admixture proportions in subsequent panels. **b,** Comparison of z-scores computed using *Twigstats* to the corresponding theoretical values shown in **a**. **c**, Admixture simulation where sources P1 and P2 split 250 or 500 generations ago and a pulse admixture event gives rise to a target population PX 50 generations ago (see **Methods** for simulation details). Admixture proportions are computed using an *f*_4_-ratio statistic and the *Twigstats* cutoff is set to twice the source split time and the rare variant cutoff is 5%. **d,** *QpAdm* simulation taken from^7,10^, as well as an adapted version where all population split times and the admixture date are divided by 5. The *Twigstats* cutoff time is chosen to be 1200 generations (top) and 600 generations (bottom).

**Supplementary Figure 4.**
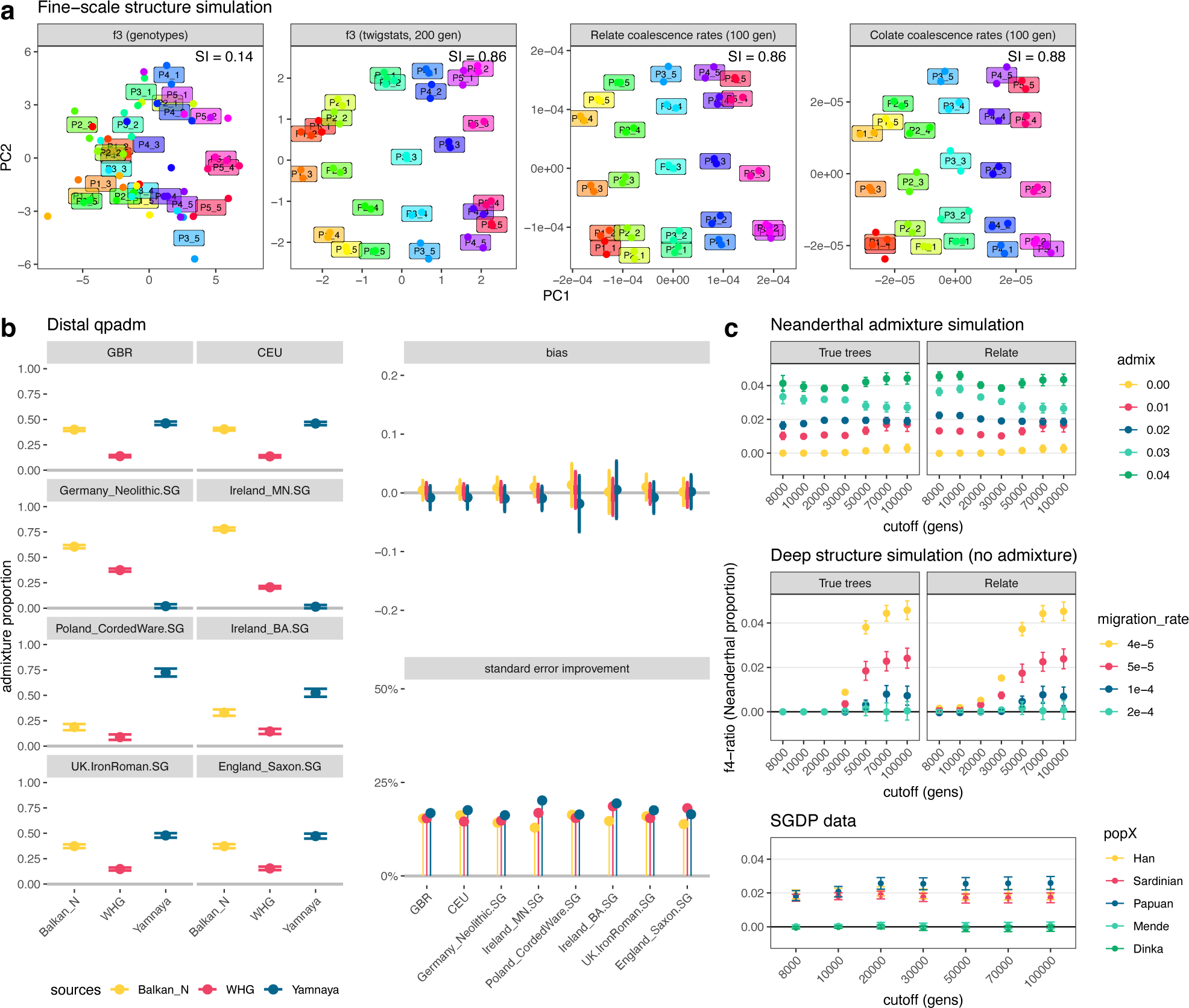
Three examples of applying *Twigstats*. **a** Fine-scale population structure simulation emulating Ref ^29^ (see Methods for simulation details). First two principal components are computed from pairwise outgroup f3 statistics on the genotypes directly and on *Relate* trees inferred from the 50 target individuals. In addition, we compute PCs on coalescence rates inferred from *Relate* trees or using *Colate^21^*. For Colate, we use a genealogy inferred only with 100 outgroup individuals, which are uncorrelated to the structure observed in the target populations and therefore emulates a situation where the genealogy is inferred on genomes that are closely related to, but separate, from the target populations of interest. Labels in plots show the average coordinates of members of that population. For each panel, we calculate a separation index as in ^29^, which we define as the proportion of individuals for which the closest individual (by the Euclidean distance in PC space) is in the same population. **b,** Admixture proportions inferred using *qpAdm* with three distal sources of Western hunter-gatherers, early European farmers, and Yamnaya Steppe people^36^. We show results for *Twigstats*-5000. Bias is measured as the difference in admixture proportions obtained from *Twigstats*-5000 and all SNPs, and we show standard errors of the latter. The standard error improvement shown is one minus the ratio of standard errors obtained from *Twigstats*-5000 and using all SNPs. **c,** Admixture proportions as a function of the *Twigstats* cutoff time computed on true or *Relate* trees for simulations emulating Neanderthal admixture or deep structure, not involving admixture, as well as for real samples in the SGDP data set ^75^.

**Supplementary Figure 5.**
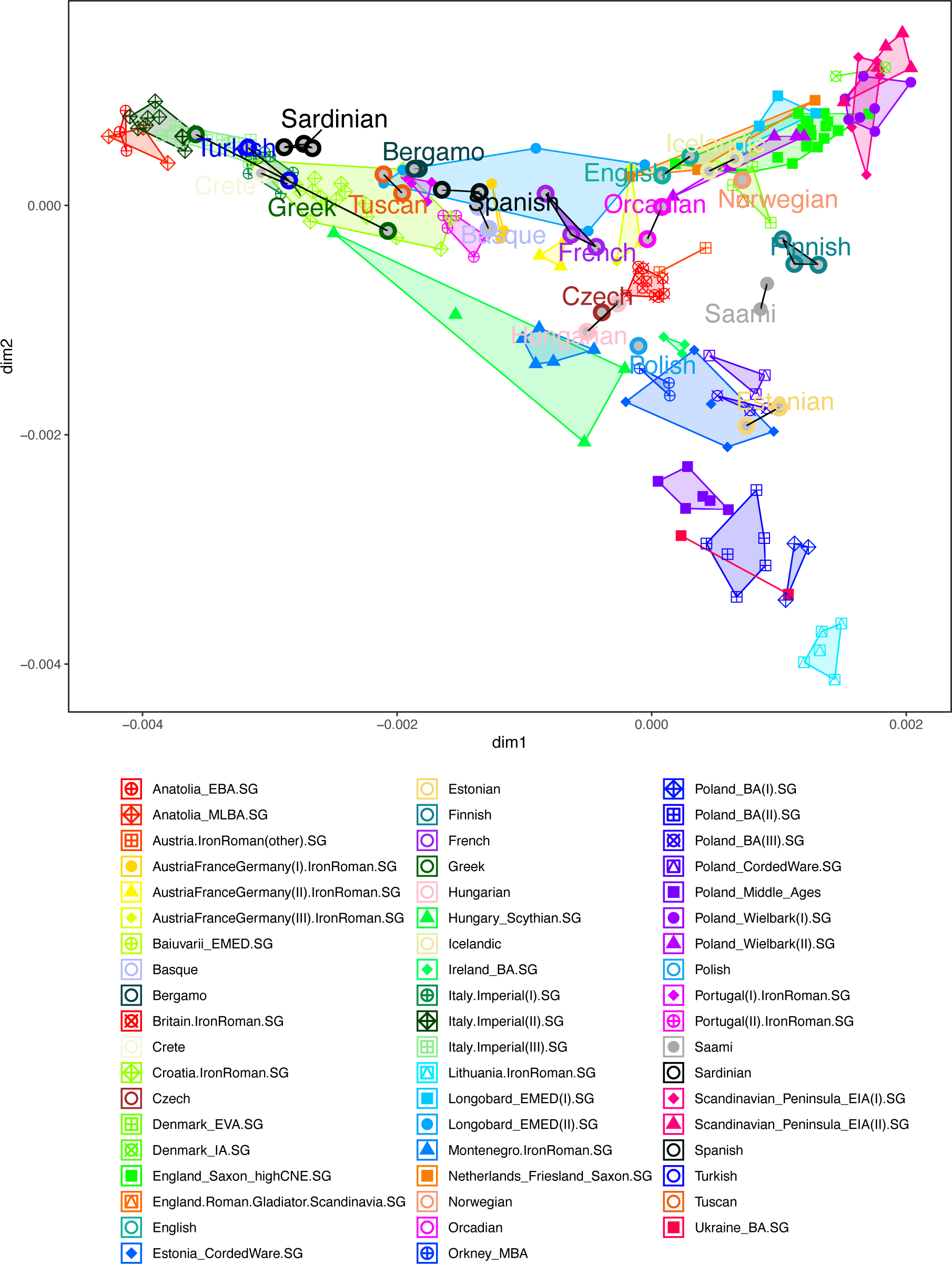
MDS of ancient and modern genomes. MDS of Iron and Roman Age groups, and modern groups in the Simons Genome Diversity Project (labelled).

**Supplementary Figure 6.**
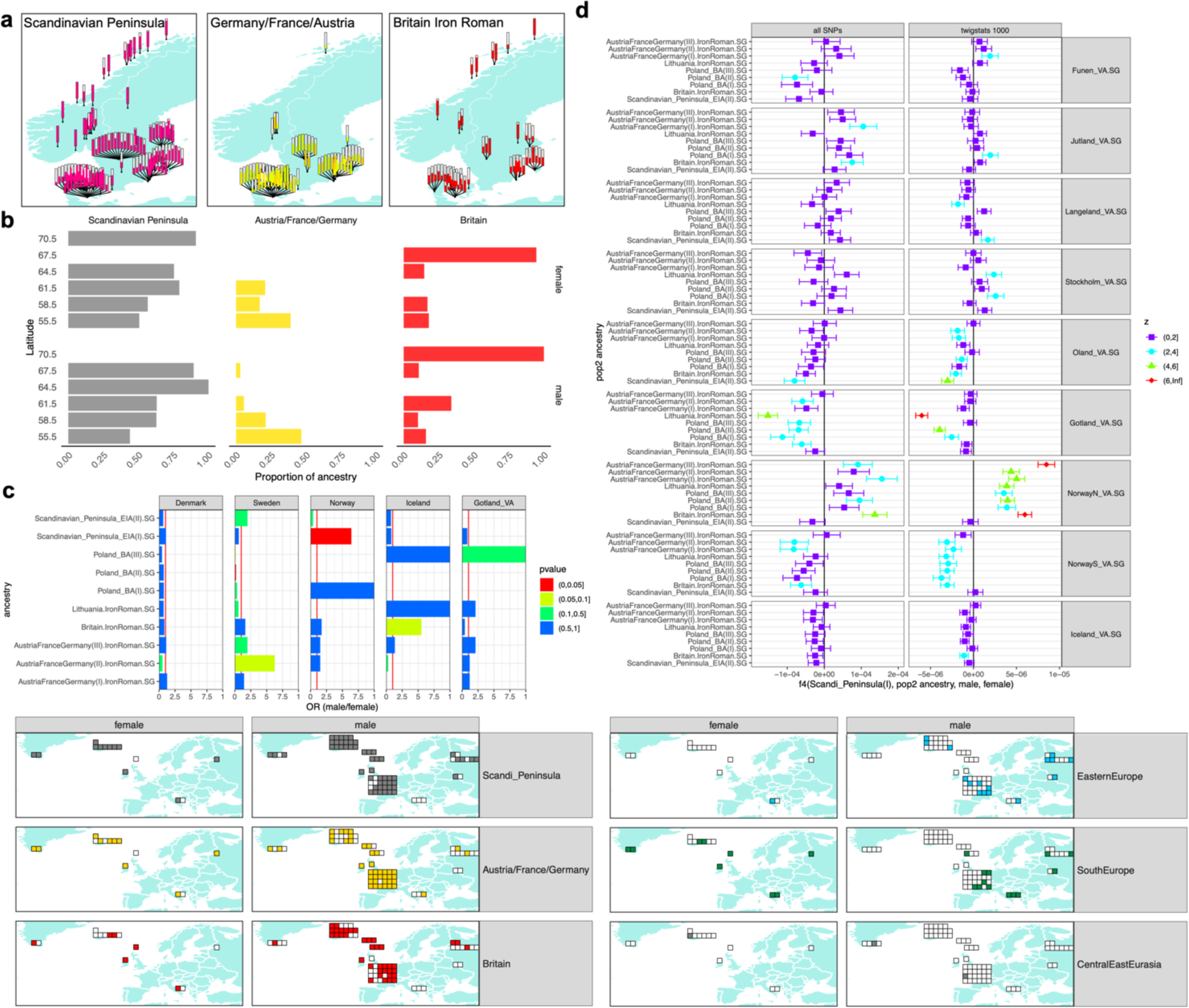
Ancestry estimates stratified by genetic sex. **a,** Map showing ancestry carried by each Scandinavian Viking age individual. **b,** Ancestry proportions across individuals grouped by Latitude and genetic sex. **c,** Odds ratio and p-values calculated using a two-sided Fisher’s exact test on the number of males and females carrying each ancestry in Viking Age Denmark, Sweden, Norway, Iceland, and Gotland. **d,** *F_4_* values of the form *f*_4_(Scandinavian_Peninsula_EIA(I), alternative source group, males in Viking group, females in Viking group) computed using all SNPs and *Twigstats*. A significantly positive value is evidence of attraction of females with pop2 or males with Scandinavian_Peninsula_EIA(I). We only show groups with at least four males and females. **e,** Map showing Farflung Viking individuals grouped by ancestry and genetic sex. In contrast to Figure 4b, for **a**, **b, c, and e**, an individual is assigned an ancestry group, if it has any accepted model (p > 0.01) where that ancestry features.

**Supplementary Figure 7.**
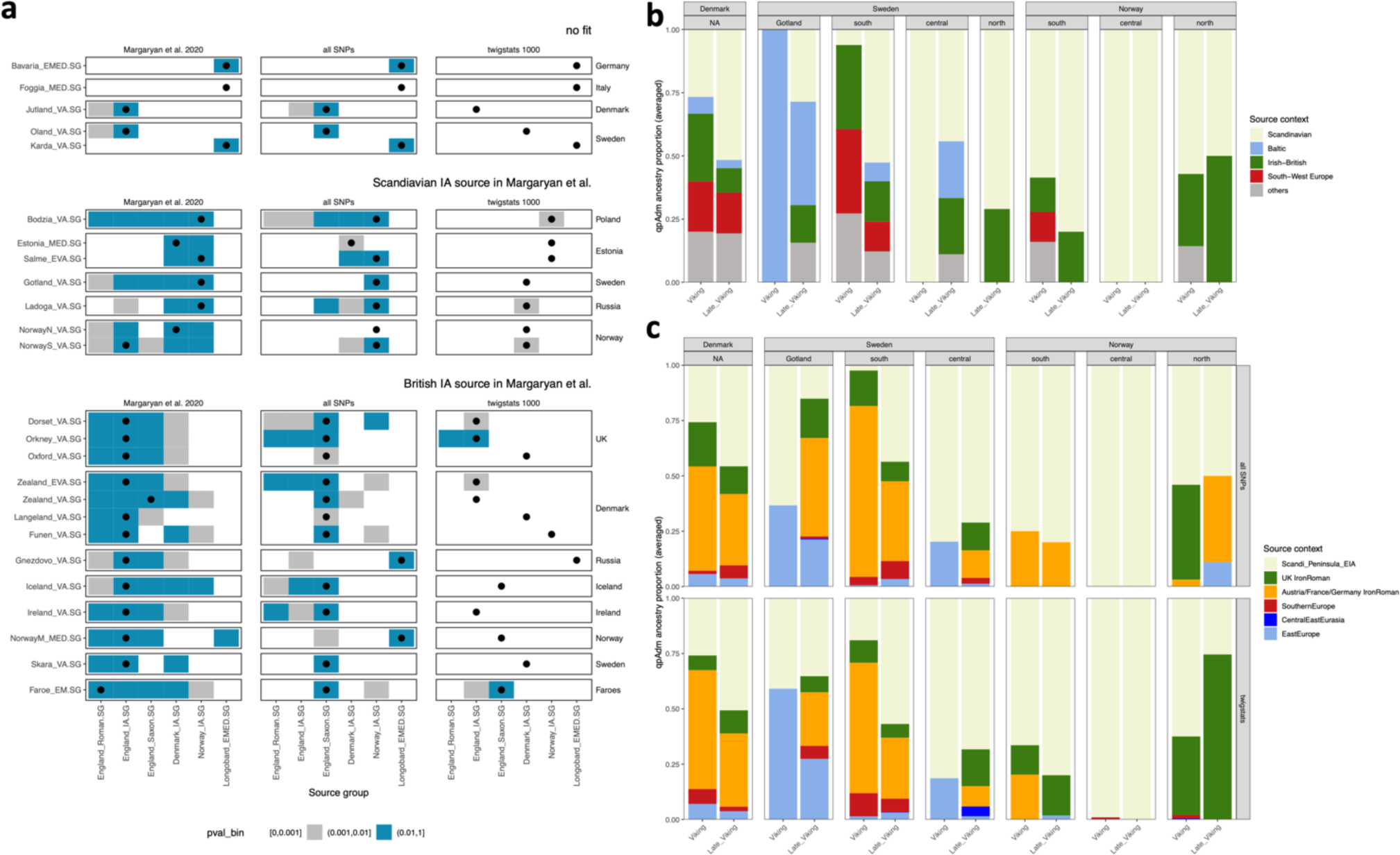
Replication previous Viking Age ancestry modelling. **a**, P-values of 1-source qpAdm models with target groups shown as rows and source groups shown as columns, replicating Supplementary Figure 5a of Ref ^13^. Left column uses p-values obtained from Ref ^13^. Middle and right column correspond to newly computed p-values in a *qpAdm* using, respectively, all SNPs and *Twigstats*-2000. Outgroups are YRI, CHB, DevilsCave_N.SG, WHG, EHG, Anatolia_N, Yamnaya, Estonia_CordedWare.SG (Supplementary Table 1). We excluded Denmark_IA.SG and England_Roman.SG from the rotational scheme as these groups overlap in ancestry with England_IA.SG and Norway_IA, respectively. Only samples with coverage exceeding 0.5 are used. For each target group, the source group with the largest p-value is shown with a black circle. **b**, *QpAdm* models of Ref ^44^ where modern populations are used as sources. As in Ref ^44^, we show ancestry proportions averaged over individuals in each group, where for each individual the model with the smallest number of sources and largest p-value is chosen. **c,** Replication using the same target samples as in **b**. We fit a maximum of two sources and choose the model with the smallest number of sources and largest p-value, requiring p > 0.05 for 1 source and p > 0.001 for 2 source models. The set of individuals used in **b** and **c** are identical and are comprised of targets with an accepted model in all SNPs and *Twigstats*-1000, removing 15 of 167 individuals. We additionally remove 17 individuals that did not have a feasible model in Ref. ^44^.

**Supplementary Figure 8.**
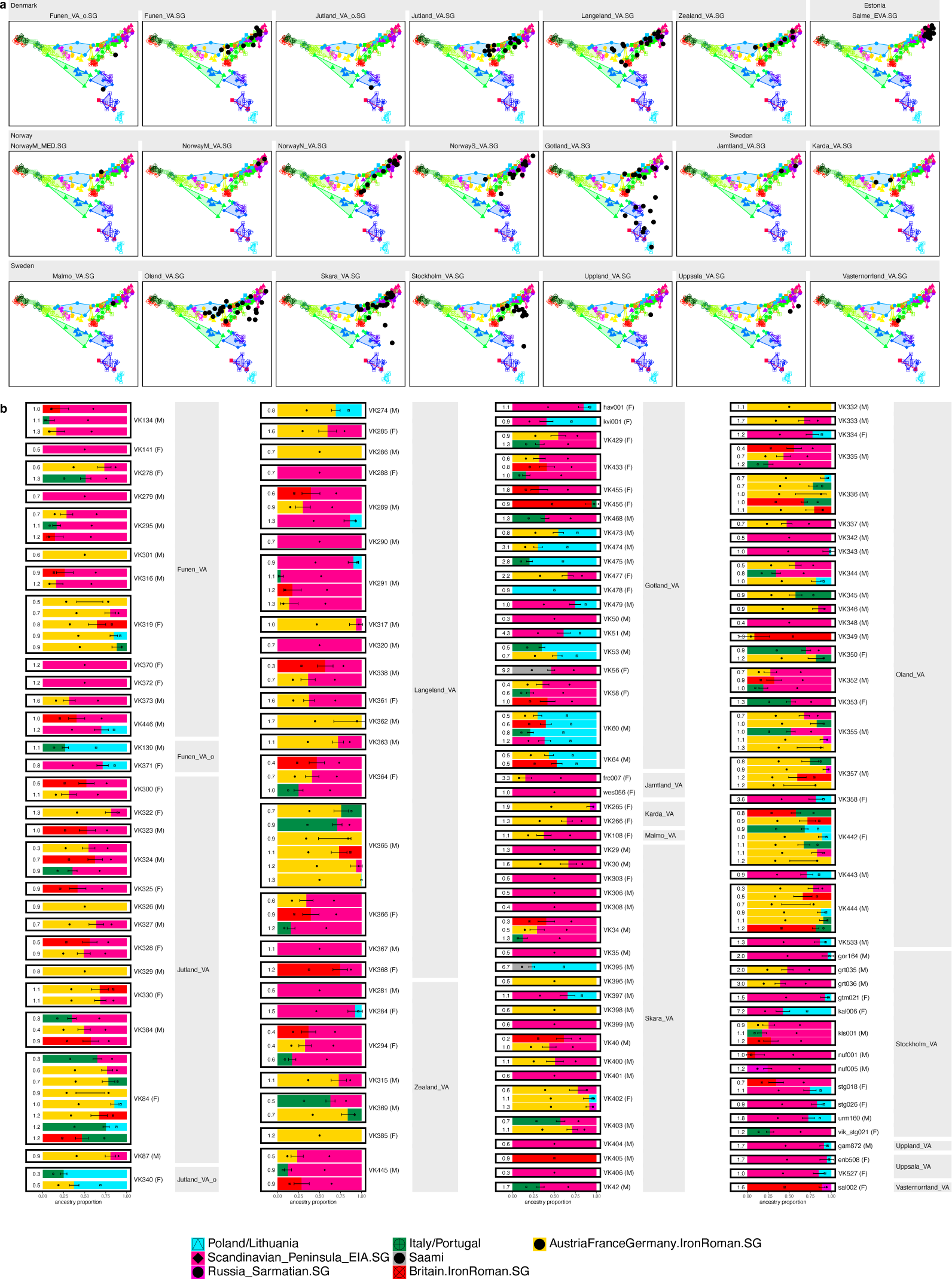
Ancestry models of Viking Age individuals in Scandinavia. **a,** MDS of each Scandinavian Viking group plotted on top of preceding Iron age and Roman individuals. **b,** All accepted qpAdm models using twigstats-1000 for every Scandinavian Viking individual in Denmark, Sweden, and Norway, computed in a rotational qpAdm with source groups identical to Figure 4. We only retain models with feasible admixture proportions, standard errors of < 0.25, and show models with 1 source and a p-value greater than 0.05 or otherwise with 2 sources and a p-value greater than 0.05. If no such models exist, we select the model with the largest p-value. The -log10 p-values are shown to the left of each model. We combine models involving related sources, if they exist, by averaging their respective admixture proportions, standard errors, and p-values.

**Supplementary Figure 9.**
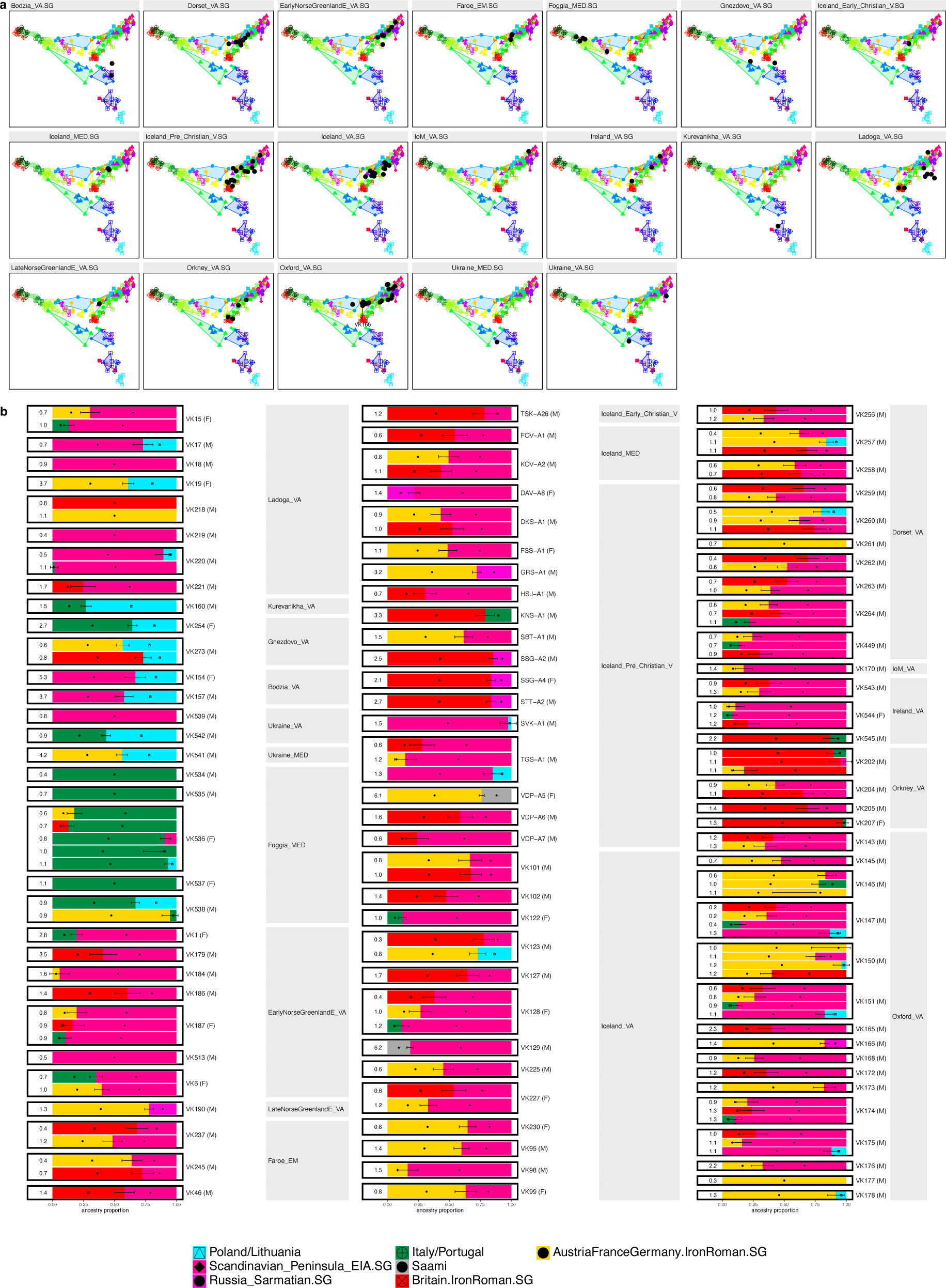
Ancestry models of farflung Viking individuals. **a,** MDS of each farflung Viking group plotted on top of preceding Iron age and Roman individuals. **b,** All accepted qpAdm models using twigstats-1000 for every non-Scandinavian Viking individual computed in a rotational qpAdm with source groups identical to Figure 4.

