## Supplementary Information for "High-resolution genomic ancestry reveals mobility in early medieval Europe"

### 1 Time-stratified $f$-statistics

Our *Twigstats* approach time-stratifies $f$-statistics by only considering lineages or mutations within a specified time period.

Specifically, our approach aims to boost statistical power of $f$-statistics for capturing recent demographic events by ascertaining recent genetic variation. Consider two populations that are recently separated. In such a case, looking backward in time, no coalescences should occur before these populations merge, and observing a lack of such recent coalescences is highly indicative of their reproductive isolation. However, since old coalescences are common, these may dominate the values of $f$-statistics or the variance of $f$-statistics across the genome, and reduce power to detect recent events. We therefore expect a statistical benefit to exclude old coalescence events that are less informative.

Computation of $f_2$-statistics from genealogies and date ascertained mutations is implemented in an R package (*Twigstats*), which is compatible with the admixtools R package (https://github.com/uqrmaie1/admixtools) [1]. The admixtools R package framework utilises the fact that f3 and f4 statistics can be calculated as functions of f2 statistics and allows us to easily access other well-used $f$-statistics-derived approaches, including *qpAdm*.

### 2 The optimal *Twigstats* cutoff

The time cutoff that minimises the standard error of computed $f$-statistics is a priori unknown but one can make an informed guess. It is useful to think of the best cutoff time when true underlying genealogies are known and $f$-statistics are calculated directly on these genealogies. As we will demonstrate here, the optimal *Twigstats* cutoff exceeds the split time of source groups involved in the admixture event by at most a factor of 1.4.

#### 2.1 Decomposing f4-statistics by how lineages coalesce back in time

Let us consider the following scenario. We consider a group $P_X$ resulting from a two-way admixture between source groups $P_1$ and $P_2$ some time $t_a$ ago, with the two source groups separating at time $t_s > t_a$ ago. For simplicity, let us assume that there is an outgroup $P_O$, which is sufficiently removed such that it does not interfere with the optimal *Twigstats* cutoff.

Let us denote by $\tau$ the *Twigstats* cutoff time. Let us assume that there are $L$ independent loci in the genome. We denote by $a_P$ the frequency of alleles in each population $P$. We are interested in computing an $f_4$-statistic of the form

$$f_4^{(\tau)}(P_O, P_2, P_X, P_1) = \frac{1}{L}\sum_{\ell=1}^{L} E_{\ell,\tau}\left[(a_O - a_2)(a_X - a_1)|T_\ell\right], \tag{1}$$

which is negative if $P_X$ carries ancestry from $P_2$ and zero if $P_X$ is unadmixed and instead clusters with $P_1$ to the exclusion of $P_2$. The expectation in Eq. (1) is taken over branches in the marginal coalescent tree $T_\ell$ at locus $\ell$ assuming a constant mutation rate. Eq. (1) is therefore a random variable that depends on the realised marginal trees at $L$ loci. We will decompose the $f_4^{(\tau)}$ statistic by considering different possible marginal trees.

We now sample one lineage from each of the four populations $P_X$, $P_1$, $P_2$, and $P_O$. Allele frequencies $a_X, a_1, a_2, a_O$ can therefore only take values 0 or 1. Let us denote by $\{X \to 2\}$ and $\{X \to 1\}$ the events that the lineage in $X$ migrates to population $P_2$ or $P_1$ back in time. These events occur with probability $\alpha$ and $1-\alpha$, respectively, corresponding to the admixture proportions. We note that migration does not imply coalescence and that a lineage that migrated to $P_2$ may not coalesce until the source split time $t_s$ and can then coalesce with $P_1$ before coalescing with $P_2$. We assume a constant effective population size throughout, and set this without loss of generality to 1.

We can now decompose our $f_4^{(\tau)}$-statistic and obtain,

$$f_4^{(\tau)} = \mathbb{1}_{\{X\to 1\}} f_4^{(\tau)}(X \to 1) + \mathbb{1}_{\{X\to 2\}} f_4^{(\tau)}(X \to 2) \tag{2}$$

Let us further denote by $\{(X,1);t\}$, $\{(X,2);t\}$, and $\{(1,2);t\}$ the disjoint events that the first coalescence of these lineages occurs between the populations shown in brackets and at time $t$, ignoring for now which population the lineage in $P_X$ migrated to. Each of these events, when realised, contribute a different amount to our $f_4$-statistic in Eq. (1).

The event $\{(X,1);t\}$ always contributes 0 to the $f_4^{(\tau)}$ statistic, the event $\{(X,2);t\}$ contributes a negative value that depends on time $t$, and $\{(1,2);t\}$ contributes a positive value that depends on time $t$. The exact value that is being contributed is computed by integrating over possible times when the coalesced lineages coalesces with the remaining third lineage, and is therefore a function of time $t$. We further note that because all effective population sizes are the same, and sample sizes in each population are identical (equalling 1), these events are equally likely if $t > t_s$, or in other words it does not matter whether the lineage of population $X$ migrated to population 1 or 2.

We use these events to further decompose the $f_4^{(\tau)}$ statistics. Starting with $f_4^{(\tau)}(X \to 1)$, a coalescence $t < t_s$ is only possible with population 1 which contributes 0, and therefore, the only non-zero contributions arise when the first coalescence is $t > t_s$ and we obtain,

$$f_4^{(\tau)}(X \to 1) = \mathbb{1}_{\{(X,2);t>t_s\}} f_4^{(\tau)}((X,2); t > t_s) + \mathbb{1}_{\{(1,2);t>t_s\}} f_4^{(\tau)}((1,2); t > t_s). \tag{3}$$

Next, we can decompose $f_4^{(\tau)}(X \to 2)$. Here, a coalescence at times $t < t_s$ with population 2 contributes negatively, and so we obtain,

$$f_4^{(\tau)}(X \to 2) = \mathbb{1}_{\{(X,2);t\le t_s\}} f_4^{(\tau)}((X,2); t \le t_s) + \mathbb{1}_{\{(X,2);t>t_s\}} f_4^{(\tau)}((X,2); t > t_s) + \mathbb{1}_{\{(1,2);t>t_s\}} f_4^{(\tau)}((1,2); t > t_s). \tag{4}$$

By substituting Eqs. (3) and (4) in Eq. (2), we obtain

$$f_4^{(\tau)} = \mathbb{1}_{\{X\to 2\}} \mathbb{1}_{\{(X,2);t\le t_s\}} f_4^{(\tau)}((X,2); t \le t_s) + \mathbb{1}_{\{(X,2);t>t_s\}} f_4^{(\tau)}((X,2); t > t_s) + \mathbb{1}_{\{(1,2);t>t_s\}} f_4^{(\tau)}((1,2); t > t_s). \tag{5}$$

#### 2.2 Expected value of the f4-statistics

We can now compute the expectation of Eq. (5). Since all coalescence rates are equal at $t > t_s$, we note that

$$f_4^{(\tau)}((X,2); t > t_s) = -f_4^{(\tau)}((1,2); t > t_s), \tag{6}$$

and because events $\{(X,2);t\}$ and $\{(1,2);t\}$ are equally likely, we get that the last two terms in Eq. (5) cancel out in expectation. The probability of the lineage in population $X$ migrating to population 2 is *alpha* and the probability of it coalescing with the lineage in population 2 before $t_s$ is given by $1 - e^{-(t_s - t_a)}$ and we therefore obtain

$$E[f_4^{(\tau)}] = P(X \to 2)P\left((X,2); t \leq t_s\right) E[f_4^{(\tau)}((X,2); t \leq t_s)] \tag{7}$$

$$= \alpha \left(1 - e^{-(t_s - t_a)}\right) E[f_4^{(\tau)}((X,2); t \leq t_s)]. \tag{8}$$

In other words, terms that on-average have a non-zero contribution on the $f_4^{(\tau)}$ statistic arise only through the combination of events $\{X \to 2\}$ and $\{(X,2); t \leq t_s\}$ and all other terms, which correspond to cases where the first coalescence occurs at times $t > t_s$, purely contribute noise to the $f_4^{(\tau)}$ statistic. The term $E[f_4^{(\tau)}((X,2); t \leq t_s)]$ corresponds to the expected number of mutations observed on a branch where $X$ and 2 coalesce.

This indicates a tradeoff: We expect a lower *Twigstats* cutoff to increase power because it reduces the chance of a coalesce $t > t_s$ from occurring, which only contribute noise. This is because lineages either coalesce before $t_s$ or, with increasing chance, don't coalesce before the *Twigstats* cutoff. On the other hand, once a coalescence $t \leq t_s$ between $X$ and 2 has occurred, which will have a non-zero contribution to the $f_4^{(\tau)}$-statistic that is maximally informative of the admixture, we now want to maximally count mutations on the resulting coalesced lineage, i.e. ideally count the entire branch until it coalesces with the remaining lineage originally in population 1. This is achieved by a larger cutoff time.

#### 2.3 Derivation of the optimal *Twigstats* cutoff

Given this trade-off, there should be an optimal choice for the *Twigstats* cutoff. To find this optimal value, we are interested in maximising

$$\frac{E[f_4^{(\tau)}]}{\sqrt{Var[f_4^{(\tau)}]}}, \tag{9}$$

which can be interpreted as a z-score, or inverse coefficient of variation. It will suffice to compute these at a single locus and the genome-wide value is obtained by multiplying by $\sqrt{L}$, assuming we have $L$ independent loci.

It will be convenient to square Eq. (9) and we obtain

$$\frac{E\left[f_4^{(\tau)}\right]^2}{Var\left[f_4^{(\tau)}\right]} = \frac{E\left[f_4^{(\tau)}\right]^2}{E\left[\left(f_4^{(\tau)}\right)^2\right] - E\left[f_4^{(\tau)}\right]^2} = \left(\frac{E\left[\left(f_4^{(\tau)}\right)^2\right]}{E\left[f_4^{(\tau)}\right]^2} - 1\right)^{-1}, \tag{10}$$

and we therefore need to compute the first two moments of our $f_4^{(\tau)}$-statistic. The first moment is given by Eq. (7). For the second moment, we use our decomposition of the $f_4^{(\tau)}$-statistic in Eq. (5). Since Eq. (5) is a sum of disjoint events, we can simply square each term. We note that the probability of an event $\{(X,2); t > t_s\}$ is given by $e^{-(t_s - t_a)}/3$, because the $P_X$ lineage is not allowed to coalesce before time $t_s$ with either $P_1$ or $P_2$ (depending on which population it migrated to) and then has to coalesce with $P_2$ which has a chance of 1/3. This probability is the same for event $\{(1,2); t > t_s\}$. We therefore obtain

$$E\left[(f_4^{(\tau)})^2\right] = \alpha\left(1 - e^{-(t_s-t_a)}\right) E\left[\left(f_4^{(\tau)}((X,2); t \le t_s)\right)^2\right] \tag{11}$$

$$+ \frac{e^{-(t_s-t_a)}}{3}\left(E\left[\left(f_4^{(\tau)}((X,2); t > t_s)\right)^2\right] + E\left[\left(f_4^{(\tau)}((1,2); t > t_s)\right)^2\right]\right) \tag{12}$$

$$= \alpha\left(1 - e^{-(t_s-t_a)}\right) E\left[\left(f_4^{(\tau)}((X,2); t \le t_s)\right)^2\right] + \frac{2e^{-(t_s-t_a)}}{3} E\left[\left(f_4^{(\tau)}((1,2); t > t_s)\right)^2\right], \tag{13}$$

where we combined terms by using that all lineages are exchangable at $t > t_s$ and therefore $E\left[\left(f_4^{(\tau)}((X,2); t > t_s)\right)^2\right] = E\left[\left(f_4^{(\tau)}((1,2); t > t_s)\right)^2\right]$.

#### 2.4 Evaluating the theoretical z-score given a *Twigstats* cutoff

We can now substitute Eqs. (7) and (11) in Eq. (10). If we can evaluate the three integrals given by

- $E\left[f_4^{(\tau)}((X,2); t \le t_s)\right]$
- $E\left[\left(f_4^{(\tau)}((X,2); t \le t_s)\right)^2\right]$
- $E\left[\left(f_4^{(\tau)}((1,2); t > t_s)\right)^2\right]$

then we can compute a theoretical z-score as a function of the *Twigstats* cutoff time and search for the optimal value. We implemented a function `theoretical_zscore` computing this theoretical z-score in our *Twigstats* package. Arguments to this function are the cutoff time (in generations), admixture date, source split time, and admixture proportion.

To evaluate the integrals, we start with $E\left[f_4^{(\tau)}((X,2); t \le t_s)\right]$. We need to integrate over times when lineages $X$ and 2 coalesce, and then, if $\tau > t_s$, also over times when the coalesced lineage coalesces with 1. We obtain

$$E\left[f_4^{(\tau)}((X,2); t \le t_s)\right] = \frac{-1}{1 - e^{-(t_s-t_a)}}\left[1_{\{\tau > t_s\}}\int_{t_a}^{t_s}\int_{t_s}^{\tau}(t_2 - t_1)e^{-(t_1-t_a)}e^{-(t_2-t_s)}dt_2\right. \tag{14}$$

$$+ (\tau - t_1)e^{-(t_1-t_a)}e^{-(\tau-t_s)}dt_1 \tag{15}$$

$$\left. + 1_{\{\tau \le t_s\}}\int_{t_a}^{\tau}(\tau - t_1)e^{-(t_1-t_a)}dt_1\right]. \tag{16}$$

The second moment $E\left[\left(f_4^{(\tau)}((X,2); t \le t_s)\right)^2\right]$ takes a similar form and we obtain

$$E\left[\left(f_4^{(\tau)}((X,2); t \le t_s)\right)^2\right] = \frac{1}{1 - e^{-(t_s-t_a)}}\left[1_{\{\tau > t_s\}}\int_{t_a}^{t_s}\int_{t_s}^{\tau}(t_2 - t_1)^2 e^{-(t_1-t_a)}e^{-(t_2-t_s)}dt_2\right. \tag{17}$$

$$+ (\tau - t_1)e^{-(t_1-t_a)}e^{-(\tau-t_s)}dt_1 \tag{18}$$

$$\left. + 1_{\{\tau \le t_s\}}\int_{t_a}^{\tau}(\tau - t_1)^2 e^{-(t_1-t_a)}dt_1\right]. \tag{19}$$

Finally, the second moment $E\left[\left(f_4^{(\tau)}((1,2); t > t_s)\right)^2\right]$ equals 0 if $\tau \le t_s$ and so is given by

$$E\left[\left(f_4^{(\tau)}((1,2); t \le t_s)\right)^2\right] = 1_{\{\tau > t_s\}}\left[\int_{t_s}^{\tau}\int_{t_1}^{\tau}(t_2 - t_1)^2 3e^{-3(t_1-t_s)}e^{-(t_2-t_1)}dt_2 dt_1\right. \tag{20}$$

$$\left. + \int_{t_s}^{\tau}(\tau - t_1)^2 3e^{-3(t_1-t_s)}e^{-(\tau-t_1)}dt_1\right] \tag{21}$$

These integrals are straight-forward to compute using integration by parts, which is implemented in the `theoretical_zscore` function in our *Twigstats* package (`https://github.com/leospeidel/twigstats`).

##### 2.5 Evaluating the optimal *Twigstats* cutoff

We use our function `theoretical_zscore` that computes the theoretical z-score of Eq. (9) to evaluate how the optimal *Twigstats* cutoff is impacted by admixture date, source split time, and admixture proportion.

The first panel in Extended Data Figure 3a shows how the theoretical z-score varies as a function of the admixture proportion. In the second panel, we observe that a younger source split time will lead to a larger fold-increase in the optimal z-score, confirming our hypothesis that the *Twigstats* ascertaiment scheme is particularly beneficial with closely related source groups. The third panel shows that the optimal *Twigstats* cutoff is greater than the source split time and scales with both the admixture proportion and source split time. The fourth panel shows that the optimal cutoff appears to be bounded by by a factor of less than 1.4 times the source split time. We observe a good agreement with the optimal *Twigstats* cutoff observed when simulating data (Extended Data Fig. 3b.)
